# Use of polygenic risk scores and other molecular markers to enhance cardiovascular risk prediction: prospective cohort study and modelling analysis

**DOI:** 10.1101/744565

**Authors:** Luanluan Sun, Lisa Pennells, Stephen Kaptoge, Christopher P Nelson, Gad Abraham, Matthew Arnold, Steven Bell, Thomas Bolton, Stephen Burgess, Frank Dudbridge, Qi Guo, Scott C Ritchie, Eleni Sofianopoulou, David Stevens, John R Thompson, Adam S Butterworth, Angela Wood, John Danesh, Nilesh J Samani, Michael Inouye, Emanuele Di Angelantonio

**Author notes:** joint first author (Luanluan Sun, Lisa Pennells, Stephen Kaptoge). joint senior author (John Danesh, Nilesh J Samani, Michael Inouye, Emanuele Di Angelantonio). Correspondence: Emanuele Di Angelantonio, Cardiovascular Epidemiology Unit, Department of Public Health and Primary Care, University of Cambridge, Cambridge CB1 8RN, UK.

## Abstract

**Background:** There is debate about the value of adding information on genetic and other molecular markers to conventional cardiovascular disease (CVD) risk predictors.

**Methods:** Using data on 306,654 individuals without a history of CVD from UK Biobank, we calculated measures of risk-discrimination and reclassification upon addition of polygenic risk scores (PRS) and a panel of 27 clinical biochemistry markers to a conventional risk prediction model (i.e., including age, sex, systolic blood pressure, smoking status, history of diabetes, total cholesterol and HDL cholesterol). We then modelled implications of initiating guideline-recommended statin therapy after the assessment of molecular markers for a UK primary-care setting.

**Findings:** The C-index was 0.710 (95% CI, 0.703-0.717) for a CVD prediction model containing conventional risk predictors alone. The C-index increased by similar amounts when adding information on PRS or biochemistry markers (0.011 and 0.014, respectively; P<0.001), and it increased still further (0.022; P<0.001) when information on both was combined. Among cases and controls, continuous net reclassification improvements were about 12% and 19%, respectively, when both PRS and biochemistry markers were added. If PRS and biochemistry markers were to be assessed in the entire primary care population aged 40-75, then it could help prevent one additional CVD event for every 893 individuals screened. By contrast, targeted assessment only among people at intermediate (i.e., 5-10%) 10-year CVD risk could help prevent one additional CVD event for every 233 individuals screened. This targeted strategy could help reclassify 16% of the intermediate-risk group to the high-risk (i.e., ≥10%) category, preventing 11% more CVD events than conventional risk prediction.

**Interpretation:** Adding information on both PRS and selected biochemistry markers moderately enhanced CVD predictive accuracy and could improve primary prevention of CVD. However, our modelling suggested that targeted assessment of molecular markers among individuals at intermediate-risk would be more efficient than blanket approaches.

## INTRODUCTION

A key strategy in the primary prevention of cardiovascular disease (CVD) is the use of risk prediction algorithms to target preventive interventions to people who may benefit from them most.^1-6^ These algorithms usually include information on conventional risk predictors, including age, sex, smoking history, history of diabetes, blood pressure, total cholesterol and high density lipoprotein (HDL) cholesterol.^1-3^ Widely practicable assay technologies have opened opportunities to enhance the accuracy of CVD risk prediction through addition of information on many molecular risk markers. For example, analysis that combines millions of genetic variants into polygenic risk scores (PRS) has shown potential to improve CVD risk prediction.^7-10^ Furthermore, several clinical biochemistry markers – both individually and in combination - have been proposed to improve prediction when added to conventional risk factors.^11-19^

Due to inter-related reasons, however, previous studies have been able to provide limited assessments.^20-22^ First, most studies have not recorded sufficient breadth of data to consider, both separately and in combination, the impact of using information on conventional risk predictors, PRS, and a panel of clinical biochemistry markers (beyond total- and HDL-cholesterol). Second, previous studies have tended to involve moderate statistical power, whereas a high degree of power is needed to make reliable comparisons using such molecular data. Third, studies have often lacked modelling of clinical implications of initiating guideline-recommended interventions (e.g., statin therapy) after the assessment of novel risk markers, yielding uncertainty about clinical utility.

Our study, therefore, aimed to address two questions. First, what is the improvement in CVD risk prediction that can be achieved when PRS and multiple clinical biochemistry markers are added to predictors used in conventional risk algorithms? We analysed 306,654 participants from UK Biobank (UKB) to assess PRS and biochemistry markers both separately and in combination. Second, what is the estimated clinical impact of using information on molecular markers for CVD prediction? We modelled the potential clinical impact and evaluated the benefit of initiating statin therapy as recommended by guidelines in a contemporary primary care population setting.

## METHODS

### Study design and overview

Our study involved several interrelated components (**Figure 1**). First, we derived separate PRSs for CHD and stroke, and identified informative clinical biochemistry markers that could enhance prediction of incident CVD outcomes. Second, we studied these molecular markers using measures of risk discrimination and reclassification to quantify their incremental predictive value on top of conventional risk predictors. Third, to estimate the potential for disease prevention, we used a separate large dataset (based on contemporary computerised records from general practices in the UK) to adapt (i.e., recalibrate) our findings to the context of a primary prevention population eligible for CVD screening. Fourth, we modelled the clinical implications of initiating statin therapy as recommended by current guidelines comparing two different scenarios: a “blanket” approach (i.e., assessment of molecular markers in all individuals eligible for CVD primary prevention) and a “targeted” approach (i.e., focusing molecular assessment only in people judged to be at intermediate 10-year risk of CVD after initial screening with conventional risk predictors alone). Fifth, to help contextualise the potential clinical gains afforded by assessing PRS or a biochemistry panel, we compared them in the same dataset with the gains afforded by assessment of C-reactive protein alone.

**Figure 1:**
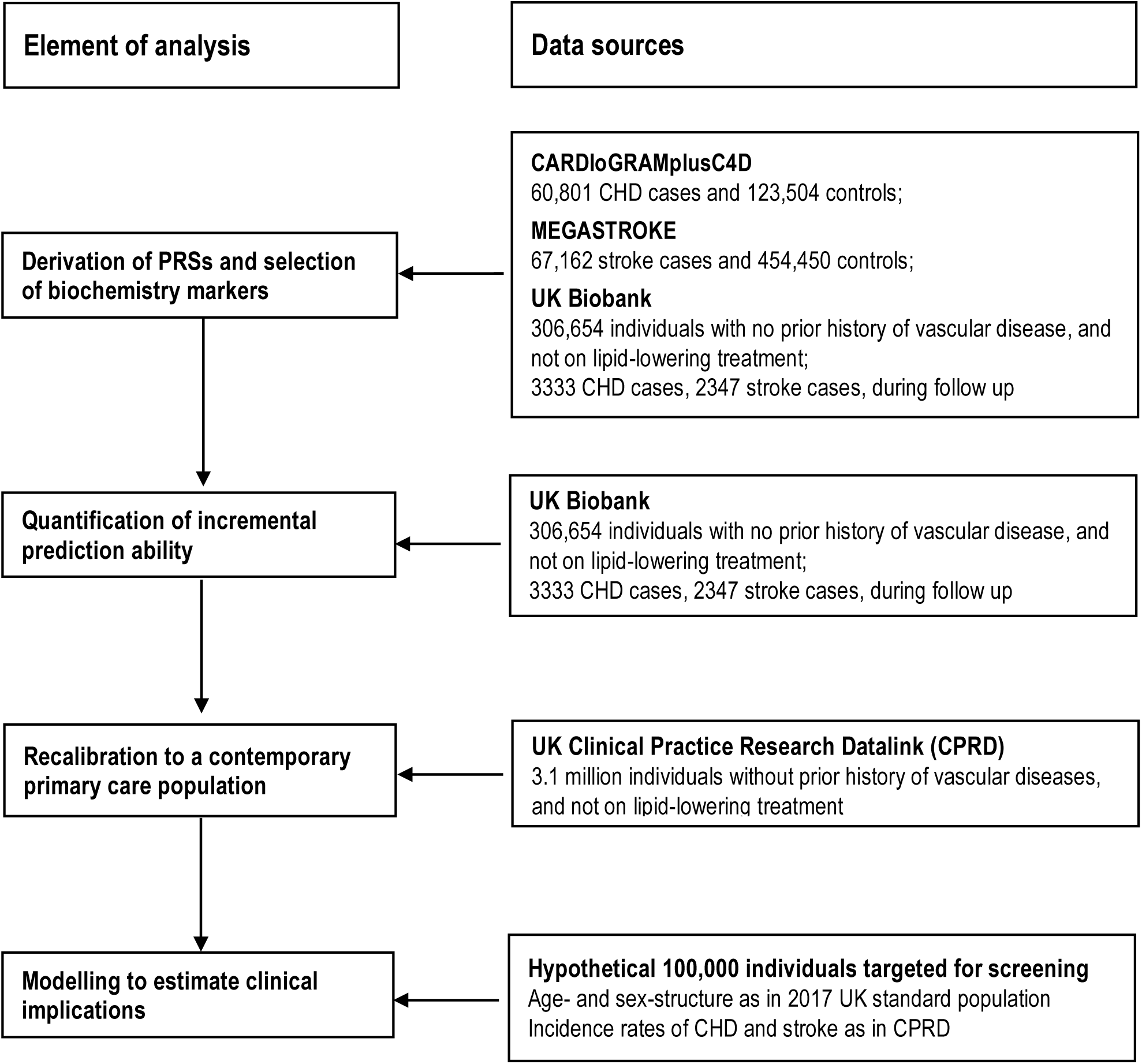
Study design. PRS, Polygenic Risk Score. CHD, Coronary Heart Disease;

### Data sources

#### UK Biobank prospective study

Details of the design, methods, and participants of UKB have been described previously.^23,24^ Briefly, participants aged 40 to 75 years identified through primary care lists were recruited across 22 assessment centres throughout the UK between 2006 and 2010. At recruitment, information was collected via a standardized questionnaire and selected physical measurements. Details of the data used from UKB are provided in the **Supplementary Appendix 1**. Data were subsequently linked to Hospital Episode Statistics (HES), as well as national death and cancer registries. HES uses *International Classification of Diseases* (ICD)–9th and 10th Revisions to record diagnosis information, and *Office of Population, Censuses and Surveys: Classification of Interventions and Procedures*, version 4 (OPCS-4) to code operative procedures. Death registries include deaths in the UK, with both primary and contributory causes of death coded in ICD-10. Genotyping was undertaken using a custom-built genome-wide array of ∼826,000 markers.^23,25^ Imputation to ∼96 million markers was subsequently carried out using the Haplotype Reference Consortium and UK10K/1000Genomes reference panels.^25^ Thirty circulating clinical biochemistry markers were measured at baseline in serum or red blood cells.^26,27^ These markers were selected for measurement in UKB for various reasons, including their relevance as established risk factors for chronic diseases, established diagnostic markers, and/or ability to reflect phenotypes not otherwise well-assessed or feasibly measured at scale.

#### UK Clinical Practice Research Datalink

To estimate the potential for disease prevention, we used data from the Clinical Practice Research Datalink (CPRD), a primary care database of anonymised medical records, with coverage of over 11.3 million patients who have opted into data linkage from 674 general practices in the UK. Individual-level data from consenting practices in the CPRD have been linked to HES and national death registry. Details of the CPRD data used and endpoint definition are provided in the **Supplementary Appendix 2**. The present analysis involved records on a random sample of all CPRD data, including 3.1 million patients. Individuals in this database should be broadly representative of the UK general population.

### Statistical analysis

Analyses included only participants of self-reported European ancestry, excluding those who: 1) had missing genotype array or clinical biochemistry marker information; 2) had prior history of vascular disease at baseline (i.e., coronary heart disease [CHD], other heart disease, stroke, transient ischaemic attack, peripheral vascular disease, angina, or cardiovascular surgery); 3) used lipid-lowering treatment at baseline; or 4) were included in the training dataset for PRS derivation (**eFigure 1**). The primary outcome was a first CVD event, defined as CHD (i.e., myocardial infarction or fatal CHD) or any stroke. Secondary outcomes included a combination of CHD, stroke, and cardiac revascularisation procedures (i.e., percutaneous transluminal coronary angioplasty [PTCA], and coronary artery bypass grafting [CABG]). Details of endpoints definitions are in **eTable 1**.

Separate PRSs for CHD and stroke were calculated as previously described,^8,28,29^ i.e., using the sum of each participant’s genome-wide genotypes weighted by corresponding genotype effect sizes (**eFigure 2**). Briefly, the PRS for CHD included 1,743,179 variants, using a meta-scoring approach to give a weighted average of three previous PRSs derived using summary statistics from up to 60,801 cases and 123,504 controls in studies from the CARDIoGRAMplusC4D Consortium.^8,28^ The PRS for stroke included 2,595,401 variants with effect sizes taken from up to 67,162 cases and 454,450 controls in the MEGASTROKE Consortium.^29^ Of the 30 circulating clinical biochemistry markers measured in UKB, 27 biomarkers were evaluated in the current analysis (**eTable 2**) because the remainder were either sex-specific markers (i.e., testosterone and oestradiol) or had >90% with missing values (i.e., rheumatoid factor). Biomarker values reported as being below or above the detectable range of assays were replaced with the minimum and maximum value reported in the available data. Distributions of continuous predictors were checked using histograms and box plots, and predictors with positively skewed distributions were natural log-transformed. To select informative clinical biochemistry markers, forward stepwise variable selection was applied using significance threshold of P <0.0001 (**eFigure 2**), yielding the following nine markers: cystatin C, lipoprotein(a), C-reactive protein, sex hormone-binding globulin, haemoglobin A1c, creatinine, albumin, gamma-glutamyltranferse, and alanine transaminase (**eTable 3**).

**Figure 2:**
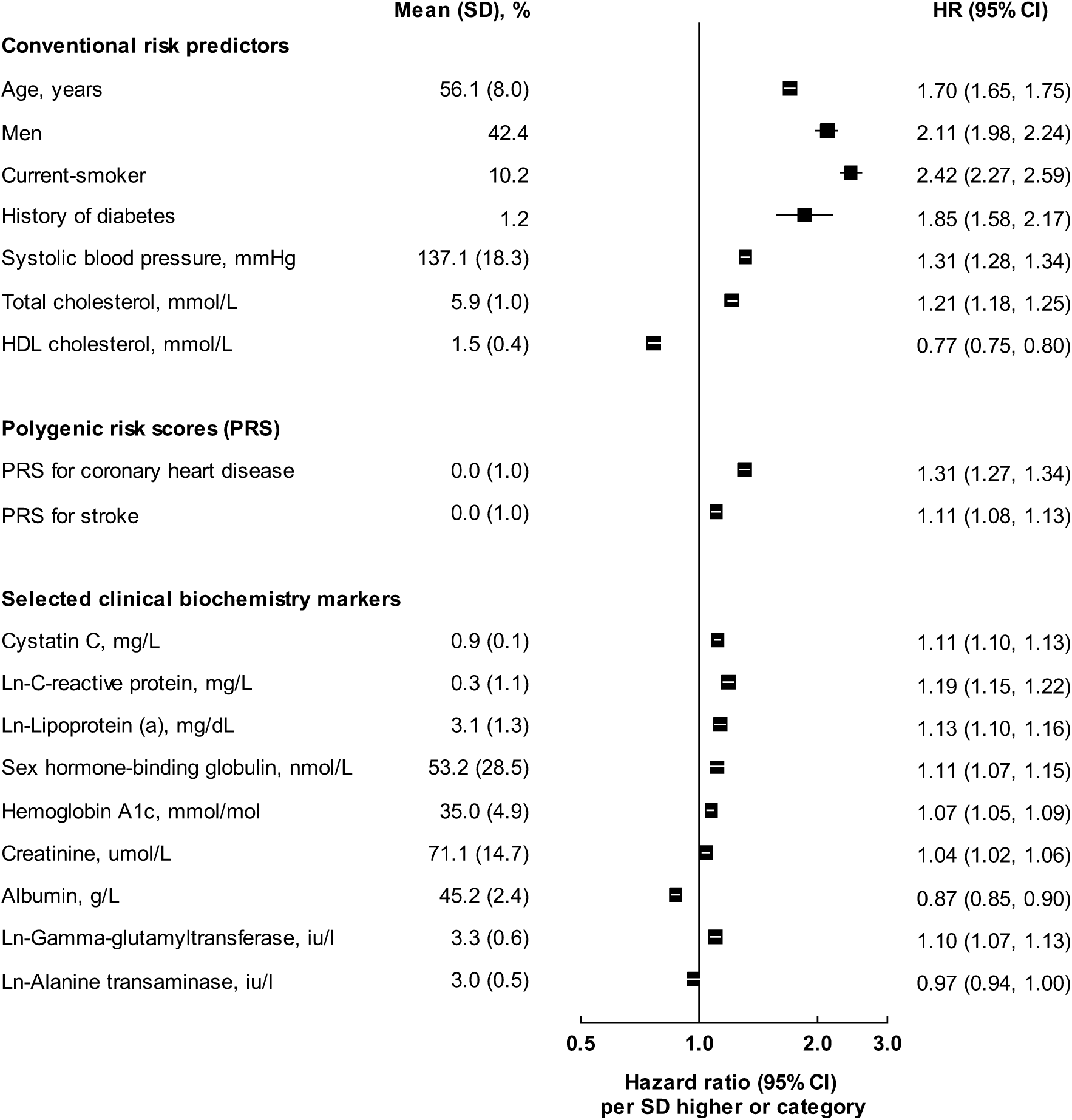
Baseline characteristics of the 306,654 participants in UK Biobank and hazard ratios for CVD adjusted for conventional risk predictors. Hazard ratios (HRs) were estimated using Cox regression, stratified by study centre and sex, and adjusted for age at baseline, smoking status, history of diabetes, systolic blood pressure, total cholesterol and HDL-cholesterol levels, where appropriate. For continuous variables, HRs are shown for each SD higher of each predictor to facilitate comparison. For categorical variables, HRs are shown for men vs. women, for patients with diabetes vs. without, for current smokers vs. others.

To quantify associations between potential risk predictors and incident outcomes, hazard ratios (HR) were calculated using Cox proportional hazards models, stratified by study centre and sex, and using time since study entry as the timescale. Outcomes were censored if a participant was lost to follow-up, died from non-CVD causes, or up to currently available end of follow-up (31 March, 2017). All predictors were entered as linear terms, after visual checking for log-linearity. No violation of the proportional hazards assumption was identified.

The incremental predictive ability of PRS and selected clinical biochemistry markers was assessed upon addition (as linear terms) to a model containing conventional CVD risk predictors, including age, sex, systolic blood pressure, smoking status, history of diabetes, total cholesterol and HDL cholesterol. Risk discrimination was assessed using Harrell’s C-index, stratified by UKB recruitment centre and sex, which estimated the probability of correctly ranking who will have an event first in a randomly selected pair of participants.^30,31^ To avoid optimism, we applied ten-fold cross-validation analyses for the main analyses. Improvements in risk prediction were also quantified using the continuous net reclassification improvement (cNRI), which summarises appropriate directional change in risk predictions for those who do and do not experience an event during follow-up (with increases in predicted risk being appropriate for cases and decreases being appropriate for non-cases).^32^

To assess the public health and clinical relevance of adding PRS and the selected biochemistry markers to conventional risk predictors, we generalised our reclassification analyses to the context of a primary prevention UK population eligible for screening (**eFigure 3**). We recalibrated risk prediction models derived in UKB to represent 10-year risks that would be expected in a UK primary care setting using CPRD data,^33^ using methods previously described.^34^

**Figure 3:**
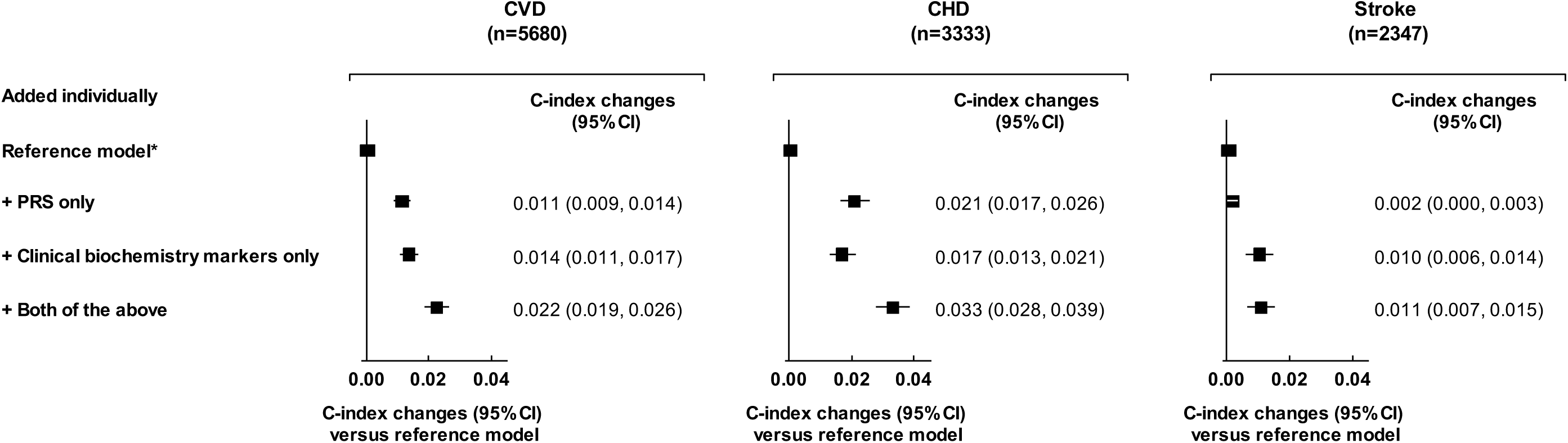
Incremental predictive ability of polygenic risk score and clinical biochemistry markers for CVD outcomes, above conventional risk predictors. CVD, cardiovascular disease; CHD, coronary heart disease; PRS, polygenic risk score; *Reference model included information on information on age at baseline, sex, smoking status, history of diabetes, systolic blood pressure, total cholesterol and HDL-cholesterol levels. PRSs for CVD included PRS for CHD and PRS for stroke as two variables. Clinical biochemistry markers were Cystatin C, C-reactive protein, Lipoprotein (a), Sex hormone-binding globulin, Hemoglobin A1c, Creatinine, Albumin, Gamma-glutamyltransferase, Alanine transaminase (Figures S2 and Table S3).

We then modelled a population of 100,000 adults aged 40-75 years, with an age and sex profile as contemporary UK population (2017 mid-year population, https://www.ons.gov.uk/), and CVD incidence rates as observed in individuals without previous CVD and not on statin treatment at registration, in the CPRD. We assumed an initial policy of statin allocation for people at ≥10% predicted 10-year risk as recommended by current National Institute for Health and Care Excellence (NICE) guidelines.^6^ We then modelled additional targeted assessment of PRS, clinical biochemistry markers, or both, among people at intermediate (5%-10% predicted 10-year risk) to estimate the potential for additional treatment allocation and case prevention, assuming statin allocation would reduce CVD risk by 20%.^35^ Details of the statistical analyses are provided in the **Supplementary Appendix 3**. Analyses were performed with PLINK 2.0,^36^ and Stata version 14, with two-sided p-values and 95% confidence intervals.

## RESULTS

### Characteristics of the study participants

Among 306,654 participants of European ancestry without a history of CVD and not on lipid-lowering treatment, the mean (SD) age was 56 (8) years, and 43% were men (**eTable 4**). During 2.6 million person-years at risk (median [5^th^,95^th^ percentile] follow-up of 8.1 [6.8-9.4] years), 3333 CHD and 2347 stroke events were recorded. Associations of PRS or biochemistry markers with CVD outcomes were approximately log-linear (**eFigures 4-5**). **Figure 2** shows the baseline characteristics of participants, as well as HRs for CVD adjusted for conventional risk predictors. HRs for CHD and stroke outcomes separately and for the composite secondary outcome (including CHD, stroke, CABG and PTCA) are presented in **eFigure 6.**

**Figure 4:**
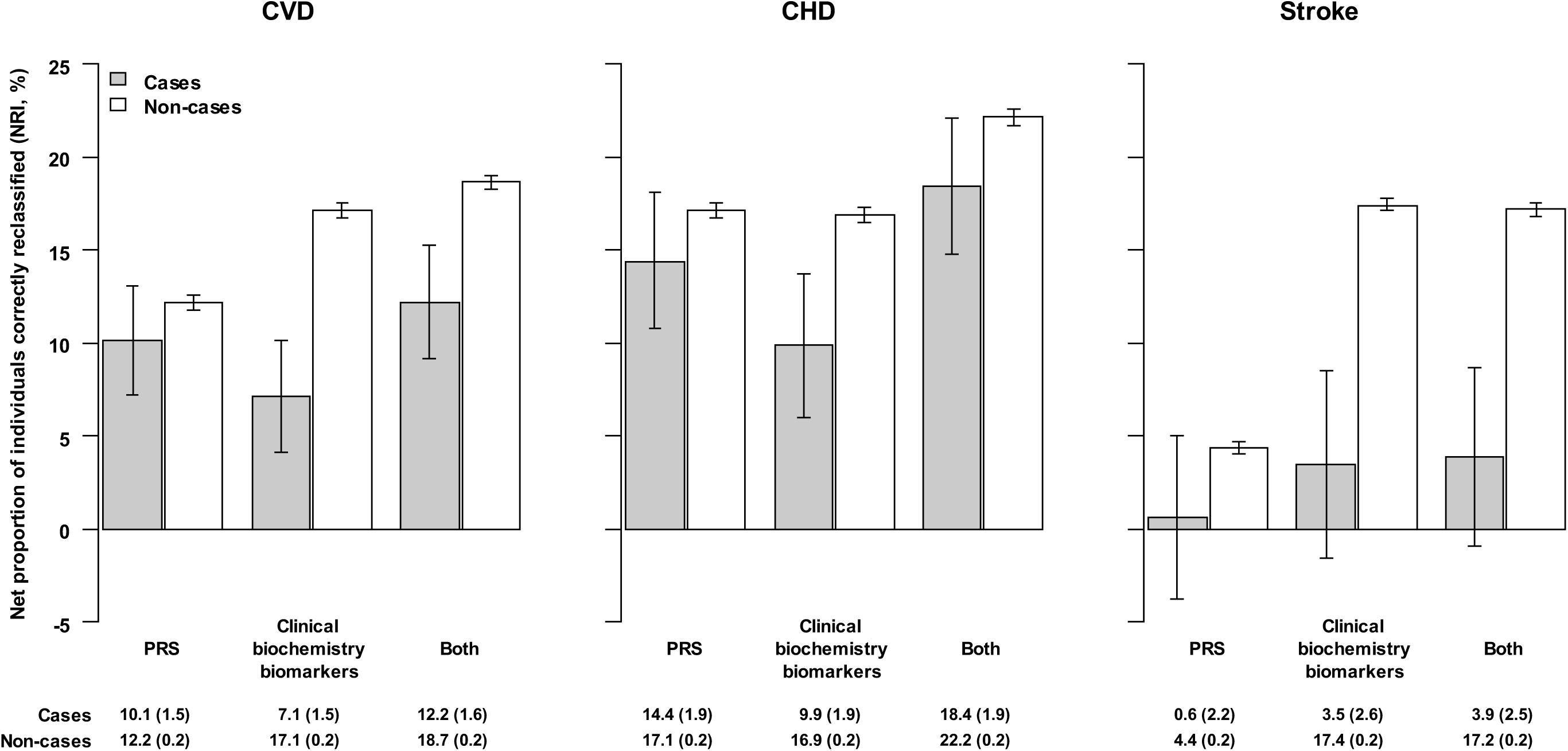
Continuous net reclassification improvement in risk prediction for CVD outcomes, with addition of information on polygenic risk score, clinical biochemistry markers, and both, to conventional risk predictors. CVD, cardiovascular disease; CHD, coronary heart disease; PRS, polygenic risk score; Conventional risk predictors included information on age at baseline, sex, smoking, systolic blood pressure, history of diabetes, total cholesterol and HDL-cholesterol levels. Clinical biochemistry markers included Cystatin C, C-reactive protein, Lipoprotein (a), Sex hormone-binding globulin, Hemoglobin A1c, Creatinine, Albumin, Gamma-glutamyltransferase, Alanine transaminase (Figures S2 and Table S3).

### Incremental value in risk prediction

The C-index was 0.710 (95% CI, 0.703-0.717) for a CVD prediction model containing conventional risk predictors alone. Addition of information on PRS or nine selected clinical biochemistry markers increased the C-index by 0.011 (0.009-0.014) and 0.014 (0.011-0.017), respectively (**Figure 3**). When both PRS and biochemistry markers were added to the model, the C-index increased by 0.022 (0.019-0.026). Incremental risk prediction afforded by PRS and/or biochemistry markers was greater than that afforded by assessment of each individual biomarker including C-reactive protein (**eFigure 7**). Among cases and controls, cNRIs for CVD were 10.1% (7.2%-13.1%) and 12.2% (11.8%-12.6%) for PRS; 7.1% (4.1%-10.1%) and 17.1% (16.7%-17.5%) for biochemistry markers; and 12.2% (9.2%-15.3%) and 18.7% (18.3%-19.0%) for both sets of markers (**Figure 4**, and **eFigure 8**). The predictive value of PRS and/or biochemistry markers was greater for CHD than for stroke outcomes (**Figures 3-4**).

The effects of adding information on PRS and biochemistry markers were similar in analyses that included: measures of body-mass index or family history of CVD or both of the preceding factors in the prediction model (**eFigure 9**); participants receiving lipid-lowering treatment at baseline (**eFigure 10**); broader definitions of CVD outcomes (i.e., CHD, stroke, PTCA and CABG; **eFigure 11**). The C-index changes with PRS were somewhat greater in men than women, but similar across age groups (**eTable 5**).

### Estimate of the potential for disease prevention

We modelled targeted assessment of PRS and biochemistry markers in a hypothetical population of 100,000 adults aged 40-75 years, assuming the current UK population age and sex structure and CVD incidence rates observed in UK primary care. Under this scenario, we estimated that, using conventional risk predictors alone, there would be 25,562 individuals classified as intermediate 10-year risk (i.e., 5%-10%) who were not already taking or eligible for statin treatment (i.e., people without a history of diabetes or CVD and with LDL-cholesterol <5.0 mmol/L; **Figure 5**). Additional assessment of PRS and clinical biochemistry markers in these individuals would reclassify 4013 individuals as high-risk (i.e., ≥10%), of whom approximately 548 would be expected to have a CVD event within 10 years. This would correspond to about 10.7% of the CVD events already classified at high risk using conventional risk predictors alone.

**Figure 5:**
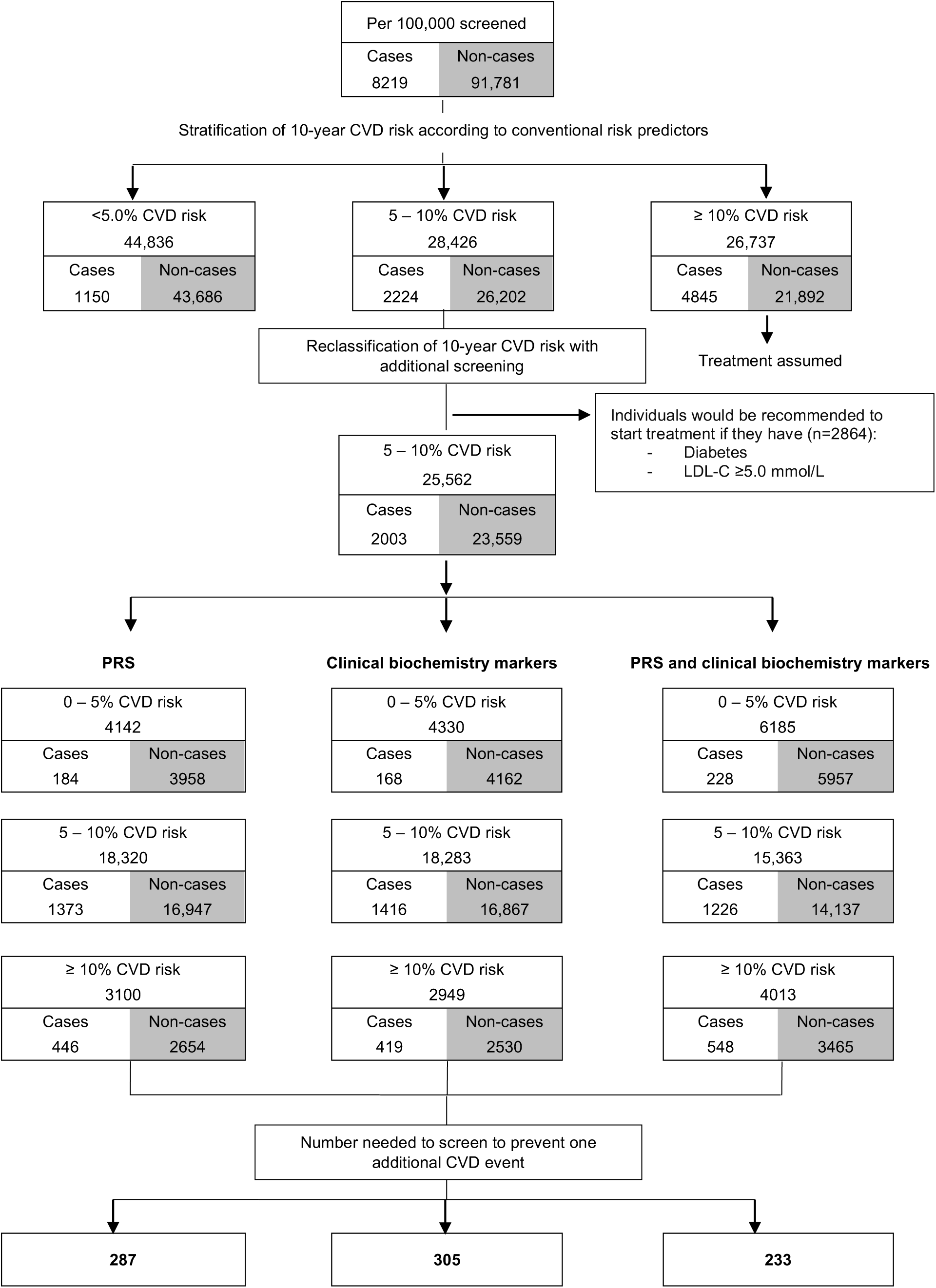
Estimated public health impact with targeted assessment of polygenic risk score, and clinical biochemistry markers among 100,000 UK adults in primary care setting

Assuming statin allocation per current guidelines (i.e., those with 10-year CVD risk ≥10%) and statin treatment conferring a 20% relative risk reduction, such targeted assessment would help prevent 110 (i.e., 548 × 0.2) events over the next 10-year period. In other words, targeted assessment of PRS and biochemistry markers in individuals at intermediate-risk for a CVD event could help prevent one additional event over 10 years for every 233 people so screened. For comparison, the corresponding number needed to screen with targeted assessment of PRS, nine selected clinical biochemistry markers, or C-reactive protein alone would be 287, 305 or 503, respectively (**Figure 5** and **eTable 6**). Similar results were observed when analysis involved cut-offs for clinical risk categories defined by the other guidelines (**eFigure 12** and **eTable 7**).

In contrast with a targeted approach, we also modelled a “blanket” strategy of assessing PRS and biochemistry markers in all adults aged 40-75 years. In this scenario, 4250 individuals would be reclassified from low- or intermediate-risk (i.e., <10%) to high-risk (i.e., ≥10%), and 5173 individuals would be reclassified from high-risk to low- or intermediate-risk, of whom approximately 561 and 443 would be expected to have a CVD event within 10 years, respectively (**eFigure 13** and **eTable 8**), suggesting the need to screen 893 people with additional assessment of PRS and biochemistry markers to help prevent one additional event over 10 years.

## DISCUSSION

Our study has yielded several findings of potential relevance to CVD risk prediction. First, our results indicate that assessment of either PRS or nine molecular markers moderately enhanced CVD prediction accuracy, affording similar gains to each other when added to conventional risk predictors. Furthermore, when information on both PRS and biochemistry markers was used in combination, the gain in predictive accuracy was largely additive, suggesting that these different sets of markers synergistically capture non-overlapping information about pathways that are uncorrelated (or weakly correlated) with conventional risk factors.

Second, our results argue in favour of targeted use of additional information on PRS and clinical biochemistry markers, rather than their “blanket” use, in screening approaches. We modelled a scenario in which PRS and biochemistry markers were assessed in a primary care setting only among individuals considered at “intermediate” risk (i.e., predicted 10-year CVD risk of 5-10%) after initial screening with conventional risk predictors alone. In a hypothetical population of 100,000 adults, we found that such targeted assessment of additional risk factors could reclassify approximately 16% (i.e., 4013/25,562) of people screened to a high-risk category (i.e., ≥10% predicted CVD risk), of whom 14% would be expected to have a CVD event within 10 years. If such an approach were to be coupled with initiation of statin therapy in accordance with guidelines, our data suggest one extra CVD outcome could be prevented over a period of 10 years for approximately every 230 people in whom both PRS and biochemistry markers are assessed. By contrast, a blanket approach in which PRS and biochemistry markers were to be assessed among all individuals relevant to a primary prevention setting would require the screening of 890 people to prevent one additional CVD outcome, i.e., a strategy that would be almost four-fold less efficient.

Third, a head-to-head comparison in our study has suggested that assessment of either PRS or a panel of biochemistry markers affords about two-fold greater predictive gains than assessment of C-reactive protein alone. This comparison might help contextualise the potential clinical gains afforded by assessing PRS and a biochemistry panel because C-reactive protein assessment alone has been recommended by some guidelines as an adjunct to CVD risk prediction.^37^ Fourth, we found that assessment of PRS and biochemistry markers could improve prediction of CHD outcomes much more than they improve prediction of stroke outcomes. Reasons for such differential gains may relate to both the greater phenotypic heterogeneity of stroke outcomes,^38-40^ and the relatively lower statistical power of previous GWAS studies of stroke,^29^ compared with CHD.^28,40^ Nevertheless, to reflect current guidelines and practice in CVD prevention, the primary outcome of our study was any first cardiovascular event (defined as fatal or nonfatal CHD or stroke).

Our study involved major strengths. We considered concomitant information on conventional CVD risk factors, PRS, and a panel of clinical biochemistry markers on more than 300,000 participants without a history of CVD at baseline. The validity of our findings was supported by the broadly concordant results we observed when using complementary measures of risk reclassification and discrimination, and by improvements in risk reclassification across the absolute risk thresholds used in different clinical guidelines. To ensure that our public health modelling was relevant to a general population, we adapted (i.e., recalibrated) the predicted risk distributions obtained from UKB to be representative of those in a primary care population.^23^

The potential limitations of this study also merit consideration. Although we used a conventional 10-year timeframe and standard clinical risk categories, we acknowledge that reclassification analyses are intrinsically sensitive to choice of follow-up interval and clinical risk categories. Furthermore, since 10 years of follow up was not available for all UKB recruitment centres, 9-year risk estimates were used in reclassification analyses. However, this limitation had minimal impact on our findings because our recalibration approach allowed reliable translation of 9-year risk estimates to 10-year risk estimates. Somewhat greater clinical impact than suggested by our analysis would be estimated if we had used less conservative modelling assumptions (e.g., use of more effective statin regimens and longer time horizons) or alternative disease outcomes (such as an exclusive focus on CHD rather than on CHD plus stroke). Conversely, our clinical models could have overestimated potential benefits of assessing PRS and clinical biochemistry markers because not all people eligible for statins will receive them or be willing, adherent, or able to take them.

A formal health economic evaluation was beyond the scope of this analysis, although we acknowledge the importance of such detailed evaluations as part of future considerations. For example, genome-wide array genotyping has a 1-time cost (approximately £40 at current prices) and can be used to calculate PRS for CVD, as well as many other chronic diseases. Future prospective studies are needed to evaluate strategies for incorporation of PRS into CVD screening, such as a “genome-first” approach that invert current “conventional risk factors first” approach to CVDs.

In summary, adding information on both PRS and selected biochemistry markers moderately enhanced CVD predictive accuracy and could improve primary prevention of CVD. However, our modelling suggested that targeted assessment of molecular markers among individuals at intermediate-risk would be more efficient than blanket approaches.

## Acknowledgements

This research has been conducted using the UK Biobank Resource under Application Number 26865. This study is based in part on data from the Clinical Practice Research Datalink obtained under licence from the UK Medicines and Healthcare products Regulatory Agency (protocol number 162RMn2). The data is provided by patients and collected by the NHS as part of their care and support. The interpretation and conclusions contained in this study are those of the author/s alone. Mortality data in CPRD was provided by the Office of National Statistics (ONS). ONS and Hospital Episode Statistics (HES) data: Copyright © (2019), re-used with the permission of The Health & Social Care Information Centre. All rights reserved. The OPCS Classification of Interventions and Procedures, codes, terms and text is Crown copyright (2016) published by Health and Social Care Information Centre, also known as NHS Digital and licensed under the Open Government Licence available at www.nationalarchives.gov.uk/doc/open-government-licence/open-government-licence.htm.

