## Supplementary Materials for "Use of polygenic risk scores and other molecular markers to enhance cardiovascular risk prediction: prospective cohort study and modelling analysis"

### Supplementary Appendix, Table of contents


| **Appendix Methods** | | **Page** |
|---|---|---|
| Appendix 1 | UK Biobank | 1 |
| Appendix 2 | UK Clinical Practice Research Datalink | 2 |
| Appendix 3 | Statistical methods used for estimating public health impact | 3 |
| **Supplementary Tables** | | |
| eTable 1 | Outcome definitions | 7 |
| eTable 2 | List of clinical biochemistry markers available in UK Biobank | 8 |
| eTable 3 | Multivariable adjusted hazard ratios for CVD of conventional risk predictors and additional selected nine biochemistry markers | 9 |
| eTable 4 | Baseline characteristics of participants in UK Biobank by sex | 10 |
| eTable 5 | Incremental predictive ability of polygenic risk score and clinical biochemistry markers, above conventional risk predictors, by sex and age at baseline | 11 |
| eTable 6 | Estimates of public health impact with targeted assessment (intermediate-risk: 5-10%) of polygenic risk score and clinical biochemistry markers, and C-reactive protein among 100,000 UK adults | 12 |
| eTable 7 | Estimates of public health impact with targeted assessment (intermediate-risk: 5-7.5%) of polygenic risk score and clinical biochemistry markers, and C-reactive protein among 100,000 UK adults | 13 |
| eTable 8 | Numerical results for estimates of public health impact by additional assessment of polygenic risk score and clinical biochemistry markers, above conventional risk predictors, among 100,000 individuals | 14 |
| **Supplementary Figures** | | |
| eFigure 1 | Exclusion criteria applied in derivation of the primary analytic dataset | 15 |
| eFigure 2 | Construction of polygenic risk score and selection of clinical biochemistry markers | 16 |
| eFigure 3 | Process of estimating public health impact | 17 |
| eFigure 4 | Shape and strength of associations of polygenic risk score with risk of CHD and stroke | 18 |
| eFigure 5a | Shape of associations of polygenic risk score and clinical biochemistry markers with risk of CVD | 19 |
| eFigure 5b | Shape of associations of polygenic risk score and clinical biochemistry markers with risk of CVD | 20 |
| eFigure 6 | Hazard ratios for CHD, stroke, and the composite CVD outcome, adjusted for conventional risk predictors | 21 |
| eFigure 7 | Incremental predictive ability of polygenic risk score and the nine selected clinical biochemistry markers for CVD outcomes, in isolation and in combination, above conventional risk predictors | 22 |
| eFigure 8 | Reclassification measures for CVD in UK Biobank by addition of polygenic risk score and clinical biochemistry markers, above conventional risk predictors | 23 |
| eFigure 9 | Incremental predictive values of polygenic risk score and clinical biochemistry markers, above conventional risk predictors, including body-mass index (BMI) or family history of CVD in the reference model | 24 |
| eFigure 10 | Incremental predictive values of polygenic risk score and clinical biochemistry markers, above conventional risk predictors, with and without excluding participants on lipid-lowering treatment | 25 |
| eFigure 11 | Incremental predictive values of polygenic risk score and clinical biochemistry markers, above conventional risk predictors, for CVD outcomes, with and without including revascularisation procedures | 26 |
| eFigure 12 | Estimates of public health impact with targeted assessment of polygenic risk score, and clinical biochemistry markers among 100,000 UK adults using AHA/ACC guideline | 27 |
| eFigure 13 | Estimates of public health impact by additional assessment of polygenic risk score and clinical biochemistry markers, above conventional risk predictors, among 100,000 individuals | 28 |
| **Supplementary References** | | 29 |

### Appendix 1. UK Biobank

Details of UK Biobank (UKB) have been described previously.[1] Briefly, over 500,000 participants aged 40-69 years were recruited during 2006-2010 in 22 geographical centres throughout the United Kingdom, covering a variety of different settings to provide socioeconomic and ethnic heterogeneity and an urban-rural mix. The assessment visit comprised electronic signed consent; a self-completed touch-screen questionnaire; brief computer-assisted interview; physical and functional measures; and collection of biological samples for all participants at recruitment for long-term storage. All participants were followed-up through linkages to routinely available national datasets, including Hospital Episode Statistics (HES) data, primary care, cancer screening data, and disease-specific registries.[2] The UK Biobank study was approved by the North West Multi-centre Research Ethics Committee, and all participants provided written informed consent to participate in the UKB.

Genotyping was undertaken using a custom-built genome-wide array of ~826,000 markers. Imputation to ~96 million markers was subsequently carried out using the Haplotype Reference Consortium and UK10K/1000Genomes reference panels. All genetic analyses utilised the UKB phase 3 release imputed genotype data.[3] Thirty circulating clinical biochemistry markers were measured in serum or red blood cells. These markers were selected for measurement in UKB for various reasons, including their reflection of established risk factors for chronic diseases, established use as diagnostic measures, and/or ability to reflect phenotypes not otherwise well-assessed or feasibly measured at scale.[4,5]

HES data available for the current analysis covered hospital admissions to NHS hospitals in England from 1996 to 2017, with the Scottish data dating back as early as 1981. HES used International Classification of Diseases (ICD)–9th and 10th Revisions to record diagnosis information, and Office of Population, Censuses and Surveys: Classification of Interventions and Procedures, version 4 (OPCS-4) to code operative procedures. Death registries provided data on deaths in the UK until 2017, with both primary and contributory causes of death coded in ICD-10. Information on smoking status (current-smoker vs. others) was obtained using the touchscreen questionnaire at baseline visit (Data-Field 20116). History of diabetes at baseline (yes vs. no) was defined using self-reported information recorded at baseline, and information collected at resurveys to populate or correct baseline data, where appropriate (Data-Fields: 2443, 4041, 10844, 20002, and 20008). Systolic blood pressure (mmHg) was measured twice on the left arm while participants were seated, using an Omron blood pressure monitor, and the mean value of the two measurements was used in the present analyses. Prior history of vascular disease was defined using self-reported information recorded at the baseline visit in UKB and updated using information on hospitalization before baseline extracted from HES. Prior vascular diseases included coronary heart disease, other heart disease, stroke, transient ischaemic stroke, peripheral vascular disease, angina, or cardiovascular surgery. Lipid-lowering status (current vs. others) was obtained via self-reported information at baseline (Data-Fields: 6177 for men and 6153 for women, respectively).

### Appendix 2. Clinical Practice Research Datalink

The Clinical Practice Research Datalink (CPRD) is an ongoing primary care database of anonymised medical records from general practitioners, with coverage of over 11.3 million patients from 674 practices in the UK (England, Wales, Scotland, and Northern Ireland), with data collected from 1987 onwards. Patients are broadly representative of the UK general population in terms of age, sex and ethnicity.[6] A subset of English practices (currently 75%, representing 58% of all UK CPRD practices) have consented to participate in the CPRD linkage scheme and have provided patient-level information. Patient-level data from consenting practices are linked via a trusted third party to other existing data sources, including HES and Office for National Statistics (ONS). Records on about 3.1 million patients were used in the present analysis.

**Disease and exposure definitions**

CVD was defined by ICD-10 codes: I21-I23 (fatal and non-fatal events), I24-I25 (fatal only) and I60-69 (fatal and non-fatal). Non-fatal hospital inpatient admissions using these codes were flagged in HES. Fatal events were flagged using death certificate information (coded using ICD-10) in the ONS data. The primary care database in CPRD is coded using medcodes (a CPRD-unique classification system), which are directly mapped to Read version 3 (CTV3) codes. Mapping of ICD-10 CVD codes to CTV3 codes was done using the NHS Digital TRUD Data Migration Toolkit;[7] the complete list of CTV3 codes generated is available on request. An individual's date of first CVD event was calculated as the earliest event flagged in any of the three sources of information. Current smokers were flagged using a code list previously reported.[8] Diabetes diagnoses were flagged by: prescriptions for anti-diabetic medications;[9] CTV3 codes indicating a diabetes diagnosis;[10] or ICD-10 codes in HES (E10-E14, G59.0, G63.2, H28.0, H36.0, Y42.3). Statin prescriptions were flagged according to a list of statin medications.[8] Full details of the code lists used are available on request.

**Calculation of CVD incidence rates**

An individual's date of entry into the CPRD cohort was defined as the latest of: 6 months after registration at GP practice contributing to CPRD; date of 30th birthday; date GP practice data quality marked as "up to standard"; 1st April 2004. The date of cohort exit was defined as the earliest of: date of de-registering at practice; date of last data upload from practice to CPRD; date of 95th birthday; date of death; or 21st Oct, 2017. Participants were assumed to be at-risk for CVD from study entry until first CVD event or censoring. The age- and sex-specific incidence rates of CVD were calculated among individuals with no record of having either CVD or a statin prescription prior to the date of study entry.

**Appendix Table. Age- and sex-specific incidence rates (per 1000 person-years) of CVD among CPRD participants (n=3,117,544)**

| Age-at-risk | Male | Female |
|---|---|---|
| 40-<45 | 2.370 | 1.042 |
| 45-<50 | 3.734 | 1.605 |
| 50-<55 | 5.569 | 2.357 |
| 55-<60 | 7.574 | 3.187 |
| 60-<65 | 10.043 | 4.495 |
| 65-<70 | 13.037 | 7.133 |
| 70-<75 | 18.763 | 11.36 |
| 75-<80 | 26.587 | 19.522 |

### Appendix 3. Statistical methods used for estimating public health impact

The general process taken to estimate the potential public health impact of using different risk models for population screening is provided in **Figure S3**, and broadly involved three pieces of information.

(1) Predicted 9-year CVD risk in UKB participants using models including conventional risk predictors, PRS and the nine selected clinical biochemistry markers.

(2) Incidence rates of CVD by sex and 5-year age-at-risk among individuals without prior history of CVD, and not on statin treatment at baseline in CPRD.

(3) UK population structure by sex and 5-year age groups in mid-2017, from UK office of national statistics.

**Calculation of 10-year CVD risk for UK Biobank participants**
Since the maximum follow-up of participants in majority of UKB centres was less than 10 years (median follow-up was 8.1 years), we opted to calculate the 9-year CVD risk for each participant as an approximation to their 10-year CVD risk, using models including the different risk predictors (e.g., conventional risk predictors, PRS, and clinical biochemistry markers). Proportional hazards assumption by centre was not violated.

**Recalibration of 10-year CVD risk for UK Biobank participants**
Given that UKB participants have been found to be, on average, healthier than the UK general population (**Appendix Figure 1**),[1] absolute risk estimates derived from UKB participants are lower than those estimated by deriving and applying risk models in a general population. This can be attributed to the higher baseline survival probability, generally lower values of risk factors, and shorter follow-up period in UKB. Crude reclassification statistics relying on clinically relevant risk thresholds, calculated within the UKB dataset are, therefore, not generalizable to a broader UK primary prevention setting. To correct for this, we adapted (i.e., recalibrated) the predicted 9-year CVD risk for each UKB participant, using incidence rates estimated in CPRD. The general recalibration process has been previously described,[11] and involves a simple rescaling of the participants' risk predictions without affecting the ability of the model to discriminate risk. For the current analysis, recalibration was undertaken separately for each model, using the following steps:

(1) In UKB, we estimated the 9-year CVD risk ($\widehat{\text{risk}}_{\text{pred},i}(9)$) for individual $i$ using a Cox model including the relevant set of risk predictors.

(2) In UKB, we calculated the mean of the predicted 9-year CVD risks for each sex and 5-year age group ($\widehat{\text{risk}}_{\text{pred,agegrp}}(9)$).

(3) In CPRD, among individuals without prior history of CVD, and not on statin treatment at registration, we calculated the incidence rates of CVD for each sex and 5-year age-at-risk group. Assuming exponential survival (i.e., constant hazard) within each 5-year age group, the expected 10-year CVD risk was estimated as follows:

$$\widehat{\text{risk}}_{\text{expected}}(10) = 1 - \exp(-IR_{\text{mid}} \times 10) \quad (1)$$

where $IR_{\text{mid}}$ is the annual incidence at the mid-point of the 10 year interval ahead, i.e., for the 40 to 44 year age-group the incidence rate for 45 to 49 years was used.

(4) The following recalibration model was fitted relating the expected risk to the means of predicted risks by age group, for each sex, with transformation applied.

$$g\left(1 - \widehat{\text{risk}}_{\text{expected}}(10)\right) = \alpha + \beta g\left(1 - \widehat{\text{risk}}_{\text{pred}}(9)\right) \quad (2)$$

where $g(.)$ is the link function $\ln(-\ln(.))$

(5) $\hat{\alpha}$ and $\hat{\beta}$ from the fitted recalibration model are then used to adjust the original 9-year risk prediction $\widehat{\text{risk}}_{\text{pred},i}(9)$ for each participant $i$ in the UKB dataset, yielding a recalibrated 10-year risk prediction $\widehat{\text{risk}}_{\text{recal},i}(10)$ using the relation:

$$\widehat{\text{risk}}_{\text{recal},i}(10) = 1 - g^{-1}\left(\hat{\alpha} + \hat{\beta} g\left(1 - \widehat{\text{risk}}_{\text{pred},i}(9)\right)\right) \quad (3)$$

**Estimation of reclassification and translation to 100,000 UK individuals**
We used the recalibrated predicted CVD risk $\widehat{\text{risk}}_{\text{recal},i}(10)$ to estimate the reclassification of individuals, between risk categories used in clinical guidelines (e.g., <5%, 5-10%, ≥10% of 10-year CVD risk).[12,13]

To express our findings in a more clinically accessible manner, we used the information observed in the reclassification tables to generalize our findings to the context of population screening. We modelled a hypothetical UK population of 100,000 individuals, with sex- and age-specific structure the same as that of the standard UK population (2017 mid-year population, https://www.ons.gov.uk/), and CVD incidence rates as observed in CPRD (**Appendix 2 Table**). We assumed that treatment with statins would reduce the risk of CVD by 20%. We assumed treatment were allocated to those: 1) estimated to be at high risk according to the recalibrated risk; 2) had history of diabetes; or 3) with LDL cholesterol levels of 190 mg/dL or greater.

**Examination of the assumptions made in recalibration process**
Our methods assume that the recalibration of predicted risks from models fitted to UKB participants gives appropriate approximation to the predicted risk distributions that would be obtained if the models were derived and applied directly in the CPRD population. By extension, we assumed that the proportions of cases and non-cases falling into clinically relevant risk categories were representative of those that would be seen in the CPRD population. Since few biomarkers and no genetic data are available in our CPRD dataset, we tested our recalibration approach using simpler prediction models involving only age, sex, smoking and diabetes status. We followed these steps:

(1) A Cox model was fitted to UKB participants, stratified by sex, and using age, smoking status and diabetes as predictors to obtain 9-year risk estimates for all UKB participants ($\widehat{\text{risk}}_{\text{pred},i}^{\text{UKB}}(9)$)

(2) Risk estimates $\widehat{\text{risk}}_{\text{pred},i}^{\text{UKB}}(9)$ were recalibrated using the CPRD incidence rates as described above, giving recalibrated 10-year risk estimates for all UKB participants ($\widehat{\text{risk}}_{\text{recal},i}^{\text{UKB}}(10)$)

(3) The same specification of Cox model in (1) was fitted to CPRD participants to obtain 10-year risk estimates for CPRD participants ($\widehat{\text{risk}}_{\text{pred},i}^{\text{CPRD}}(10)$)

(4) The distributions of $\widehat{\text{risk}}_{\text{recal},i}^{\text{UKB}}(10)$ and $\widehat{\text{risk}}_{\text{pred},i}^{\text{CPRD}}(10)$ were compared to confirm that the former is a reasonable approximation to the predicted risk distribution we would expect to see in the wider CPRD population (**Appendix 3 Figure 2**)

Good agreement between age-specific risk predictions in CRPD ($\widehat{\text{risk}}_{\text{pred},i}^{\text{CPRD}}(10)$) and the medians from risk predictions in UKB after recalibration ($\widehat{\text{risk}}_{\text{recal},i}^{\text{UKB}}(10)$) indicated that the predicted risk distribution after recalibration in UKB was representative of the predicted risk distribution in a general primary care setting.

**Appendix 3 Figure 1. Comparison of sex-specific 5-year age-at-risk incidence rates of CVD between UKB and CPRD participants**

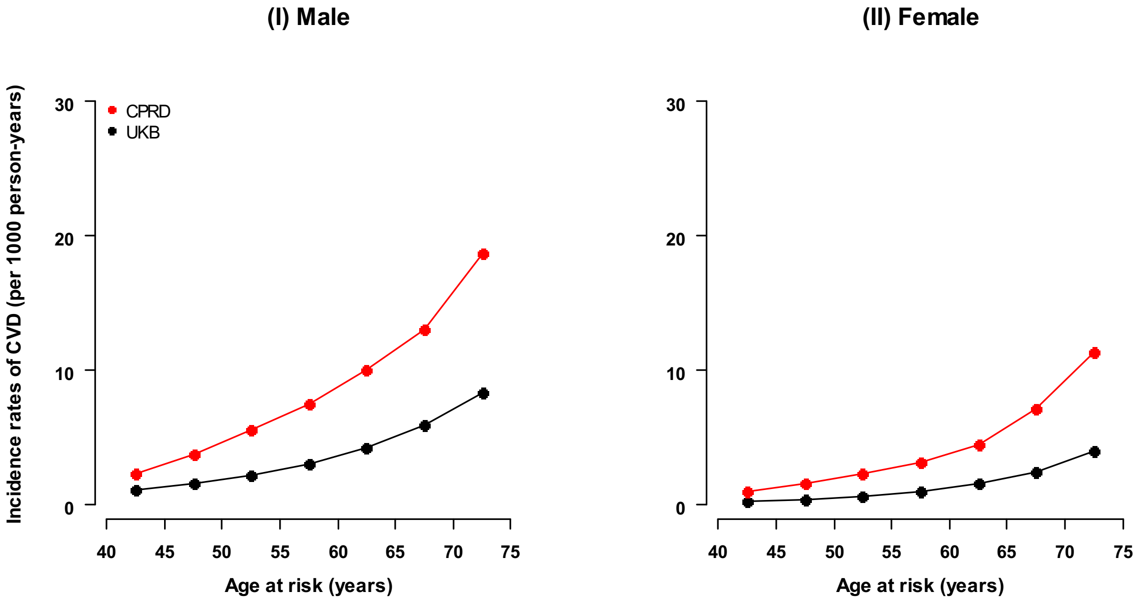


CVD, cardiovascular disease, including myocardial infarction, fatal coronary heart disease, and any stroke in both UKB and CPRD.

**Appendix 3 Figure 2: Comparison of the predicted risk in UKB after recalibration and the predicted risk in CPRD, by sex and case status at 9 years since recruitment**

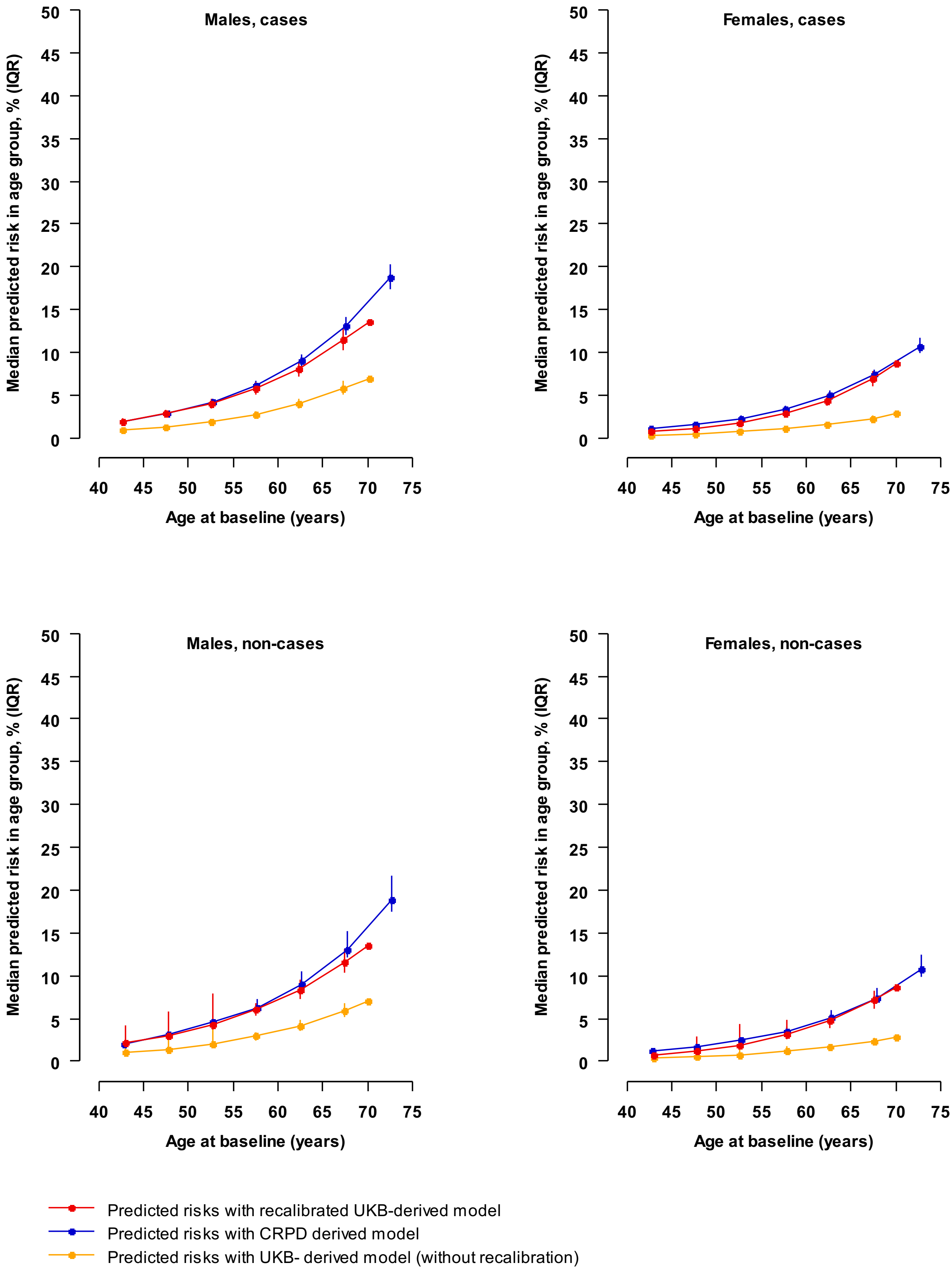

**eTable 1: Outcome definitions**

| | ICD-10 codes | OPCS-4 codes |
|---|---|---|
| ***Primary outcome*** | | |
| Cardiovascular disease (CVD) | I21-I23; fatal I24-I25, I60-69 | |
| ***Secondary outcomes*** | | |
| Coronary heart disease (CHD) | I21-I23; fatal I24-I25 | |
| Stroke | I60-69 | |
| CVD and cardiac revascularisations | I21-I23; fatal I24-I25; I60-69 | K40-K46, K49, K50.1, 50.2, K50.4, or K75 |

**eTable 2: List of clinical biochemistry markers available in UK Biobank**

| Clinical biochemistry markers | Used in the current analyses |
|---|---|
| ***Measured in serum sample*** | |
| Albumin | ✔ |
| Alkaline phosphatase | ✔ |
| Alanine aminotransferase | ✔ |
| Apolipoprotein AI | ✔ |
| Apolipoprotein B | ✔ |
| Aspartate aminotransferase | ✔ |
| Direct bilirubin | ✔ |
| Calcium | ✔ |
| Total cholesterol** | ✔ |
| Creatinine | ✔ |
| C-reactive protein | ✔ |
| Cystatin C | ✔ |
| Gamma glutamyltransferase | ✔ |
| Glucose | ✔ |
| HDL cholesterol** | ✔ |
| Insulin-like growth factor 1 | ✔ |
| Lipoprotein A | ✔ |
| LDL cholesterol | ✔ |
| Oestradiol† | |
| Phosphate | ✔ |
| Rheumatoid factor* | |
| Sex hormone-binding globulin | ✔ |
| Total bilirubin | ✔ |
| Testosterone† | |
| Total protein | ✔ |
| Triglycerides | ✔ |
| Urate | ✔ |
| Urea | ✔ |
| Vitamin D | ✔ |
| ***Measured in red blood cells*** | |
| Glycated haemoglobin | ✔ |

* Markers with >90% of missing values (i.e., rheumatoid factor), and † sex-specific markers (i.e., testosterone and oestradiol) were excluded from the following analyses.
** Total cholesterol and HDL cholesterol were included in the conventional risk predictor model.

**eTable 3: Multivariable adjusted hazard ratios for CVD of conventional risk predictors and additional selected nine biochemistry markers**

| | HR (95% CI) | Z-score |
|---|---|---|
| ***Conventional risk predictors*** | | |
| Age at baseline, year | 1.51 (1.46, 1.56) | 25 |
| Systolic blood pressure, mmHg | 1.30 (1.27, 1.34) | 20 |
| Current-smoker vs. others | 1.98 (1.85, 2.12) | 20 |
| Diabetes (Yes vs. No) | 1.42 (1.20, 1.69) | 4 |
| Total cholesterol, mmol/L | 1.20 (1.17, 1.23) | 13 |
| HDL cholesterol, mmol/L | 0.80 (0.78, 0.83) | -12 |
| **Selected nine clinical biochemistry markers** | | |
| Cystatin C, mg/L | 1.18 (1.15, 1.21) | 14 |
| Ln-C-reactive protein, mg/L | 1.10 (1.07, 1.13) | 6 |
| Ln-Lipoprotein (a), nmol/L | 1.13 (1.10, 1.16) | 9 |
| Sex hormone-binding globulin, nmol/L | 1.13 (1.09, 1.16) | 7 |
| Hemoglobin A1c, mmol/mol | 1.07 (1.05, 1.09) | 6 |
| Creatinine, umol/L | 0.92 (0.90, 0.95) | -6 |
| Albumin, g/L | 0.93 (0.90, 0.95) | -5 |
| Ln-Gamma-glutamyltransferase, iu/l | 1.12 (1.09, 1.16) | 7 |
| Ln-Alanine transaminase, iu/l | 0.91 (0.88, 0.94) | -5 |

Hazard ratios (HR) are reported per 1 SD higher for continuous predictors, stratified by recruitment centre and sex, and adjusted for all other predictors.

### eTable 4: Baseline characteristics of participants in UK Biobank by sex

| Baseline characteristics | Male | Female | Total |
|---|---|---|---|
| No. of participants | 131,881 | 174,773 | 306,654 |
| Age, years | 56 (8) | 56 (8) | 56 (8) |
| Median follow-up (5th – 95th range), years | 8.1 (6.5 to 9.3) | 8.2 (6.8 to 9.4) | 8.1 (6.8 to 9.4) |
| **Conventional risk factors** | | | |
| Current-smoker, % | 12 | 9 | 10 |
| Diabetes, % | 1.7 | 0.8 | 1.2 |
| Systolic blood pressure, mmHg | 140 (17) | 134 (19) | 137 (19) |
| Total cholesterol, mmol/L | 5.8 (1.0) | 6.0 (1.1) | 5.9 (1.1) |
| HDL cholesterol, mmol/L | 1.3 (0.3) | 1.6 (0.4) | 1.5 (0.4) |
| **Lipoprotein-related biomarkers** | | | |
| LDL cholesterol, mmol/L | 3.7 (0.8) | 3.7 (0.8) | 3.7 (0.8) |
| Apolipoprotein B, g/L | 1.1 (0.2) | 1.1 (0.2) | 1.1 (0.2) |
| Apolipoprotein A, g/L | 1.4 (0.2) | 1.6 (0.3) | 1.6 (0.3) |
| Ln-Triglycerides, mmol/L | 0.5 (0.5) | 0.3 (0.5) | 0.4 (0.5) |
| Ln-Lipoprotein(a), nmol/L | 3.0 (1.3) | 3.2 (1.3) | 3.1 (1.3) |
| **Inflammatory biomarkers** | | | |
| Ln-C-reactive protein, mg/L | 0.3 (1.0) | 0.3 (1.1) | 0.3 (1.1) |
| **Liver function biomarkers** | | | |
| Albumin, g/L | 46 (2) | 45 (2) | 45 (2) |
| Gamma-glutamyltransferase, iu/l | 3.6 (0.6) | 3.1 (0.6) | 3.3 (0.6) |
| Ln-Alkaline phosphatase, iu/l | 4.4 (0.3) | 4.4 (0.3) | 4.4 (0.3) |
| Ln-Alanine transaminase, iu/l | 3.2 (0.4) | 2.9 (0.4) | 3.0 (0.5) |
| Ln-Aspartate transaminase, iu/l | 3.3 (0.3) | 3.1 (0.6) | 3.2 (0.3) |
| Ln-Direct bilirubin, umol/L | 0.6 (0.4) | 0.3 (0.3) | 0.4 (0.4) |
| Ln-Total bilirubin, umol/L | 2.3 (0.4) | 2.0 (0.4) | 2.1 (0.4) |
| **Renal function biomarkers** | | | |
| Creatinine, umol/L | 81 (14) | 64 (11) | 71 (15) |
| Cystatin C, mg/L | 0.9 (0.1) | 0.9 (0.1) | 0.9 (0.1) |
| Urate, umol/L | 351 (69) | 266 (63) | 303 (78) |
| Urea, mmol/L | 5.5 (1.3) | 5.2 (1.2) | 5.3 (1.3) |
| **Glycaemia biomarkers** | | | |
| Haemoglobin A1c, mmol/mol | 35 (5) | 35 (4) | 35 (5) |
| Glucose, mmol/L | 5.0 (1.0) | 5.0 (0.8) | 5.0 (0.8) |
| **Others** | | | |
| Calcium, mmol/L | 2.4 (0.1) | 2.4 (0.1) | 2.4 (0.1) |
| Vitamin D, mmol/L | 49 (21) | 49 (20) | 49 (20) |
| Sex hormone-binding globulin, nmol/L | 40 (16) | 63 (32) | 53 (28) |
| Total protein (g/L) | 72 (4) | 72 (4) | 72 (4) |
| Phosphate (mmol/L) | 1.1 (0.2) | 1.2 (0.1) | 1.2 (0.2) |
| Insulin-growth factor – 1 (nmol/L) | 22 (5) | 21 (6) | 22 (6) |

Values were shown for mean (SD), stratified by centre, unless stated otherwise.

**eTable 5: Incremental predictive ability of polygenic risk score and clinical biochemistry markers, above conventional risk predictors, by sex and age at baseline**

| Subgroup | No. of Events / Total | C-index | C-index changes vs. Reference model | | |
|---|---|---|---|---|---|
| | | Reference model | + PRS only | + Clinical biochemistry markers only | + Both of the above |
| **Sex** | | | | | |
| Male | 3764 / 131,881 | 0.699 (0.690, 0.707) | 0.017 (0.014, 0.021) | 0.016 (0.013, 0.019) | 0.029 (0.025, 0.034) |
| Female | 1916 / 174,773 | 0.726 (0.714, 0.738) | 0.003 (-0.001, 0.007) | 0.011 (0.006, 0.015) | 0.013 (0.007, 0.018) |
| | | | $\chi_1$=28.6; $P_{heter}$<0.0001 | $\chi_1$=3.5; $P_{heter}$=0.063 | $\chi_1$=20.4; $P_{heter}$ <0.0001 |
| **Age at baseline, years** | | | | | |
| <55 | 1330 / 134,614 | 0.703 (0.688, 0.718) | 0.016 (0.009, 0.023) | 0.012 (0.005, 0.019) | 0.025 (0.016, 0.033) |
| 55-<65 | 2647 / 126,289 | 0.646 (0.635, 0.658) | 0.015 (0.010, 0.021) | 0.019 (0.013, 0.025) | 0.030 (0.023, 0.037) |
| >=65 | 1703 / 45,751 | 0.614 (0.600, 0.629) | 0.021 (0.013, 0.030) | 0.025 (0.016, 0.034) | 0.042 (0.032, 0.053) |
| | | | $\chi_2$=1.5; $P_{heter}$=0.47 | $\chi_2$=5.4; $P_{heter}$=0.066 | $\chi_2$=6.6; $P_{heter}$=0.037 |

Reference model included information on conventional risk predictors, i.e., age at baseline, sex, smoking status, history of diabetes, systolic blood pressure, total cholesterol and HDL-cholesterol levels. Prediction model was developed using Cox regression for all participants, stratified by study centre and sex, adjusted for conventional risk predictors, where appropriate.

**eTable 6: Estimates of public health impact with targeted assessment (intermediate-risk: 5-10%) of polygenic risk score and clinical biochemistry markers, and C-reactive protein among 100,000 UK adults**

| | **Before recalibration** | | **After recalibration** | |
|---|---|---|---|---|
| | **0-10%** | **5-10%** | **0-10%** | **5-10%** |
| ***Additional cases identified in addition to conventional risk predictors (%)*** | | | | |
| PRS only | 15.2 | 15.0 | 8.7 | 8.7 |
| Biomarkers only | 22.7 | 21.8 | 8.4 | 8.2 |
| Both of the above | 31.1 | 29.3 | 11.0 | 10.7 |
| CRP only | 9.2 | 9.2 | 5.0 | 5.0 |
| ***Number needed to screen per event prevented in addition to conventional risk predictors*** | | | | |
| *Number to screen* | *89,387* | *8871* | *68,829* | *25,562* |
| PRS only | 1832 | 183 | 768 | 287 |
| Biomarkers only | 1221 | 127 | 804 | 305 |
| Both of the above | 892 | 94 | 613 | 233 |
| CRP only | 3020 | 300 | 1355 | 503 |

PRS, polygenic risk score; CRP, C-reactive protein; Conventional risk predictors included information on age at baseline, sex, smoking, systolic blood pressure, history of diabetes, total cholesterol and HDL-cholesterol levels. The predicted 10-year cardiovascular risk categories used 2014 NICE guideline. Estimates of public health impact for a hypothetical population of 100,000 individuals (40-75 years) were based on: 1) sex- and age-specific (5-year) profile of a standard UK population (2017 mid-year population, https://www.ons.gov.uk/); 2) sex-specific 5-year age-at-risk incidence rates of CVD in CPRD, among individuals without prior history of CVD, and not on statin treatment at baseline; 3) estimates for public health impact were shown, respectively, before and after recalibration.

**eTable 7: Estimates of public health impact with targeted assessment (intermediate-risk: 5-7.5%) of polygenic risk score and clinical biochemistry markers, and C-reactive protein among 100,000 UK adults**

| | Before recalibration | | After recalibration | |
|---|---|---|---|---|
| | 0-7.5% | 5-7.5% | 0-7.5% | 5-7.5% |
| ***Additional cases identified in addition to conventional risk predictors (%)*** | | | | |
| PRS only | 12.4 | 11.4 | 4.4 | 4.1 |
| Biomarkers only | 20.4 | 18.2 | 5.7 | 5.2 |
| Both of the above | 25.8 | 17.8 | 7.1 | 6.3 |
| CRP only | 10.0 | 10.0 | 2.6 | 2.6 |
| ***Additional number screened per event prevented*** | | | | |
| *Number to screen* | *87,023* | *6505* | *58,455* | *15,187* |
| PRS only | 1667 | 136 | 1079 | 299 |
| Biomarkers only | 1007 | 85 | 835 | 238 |
| Both of the above | 800 | 87 | 664 | 197 |
| CRP only | 2052 | 153 | 1804 | 472 |

PRS, polygenic risk score; CRP, C-reactive protein; Conventional risk predictors included information on age at baseline, sex, smoking, systolic blood pressure, history of diabetes, total cholesterol and HDL-cholesterol levels. The predicted 10-year cardiovascular risk categories used 2019 AHA/ACC guideline. Estimates of public health impact for a hypothetical population of 100,000 individuals (40-75 years) were based on: 1) sex- and age-specific (5-year) profile of a standard UK population (2017 mid-year population, https://www.ons.gov.uk/); 2) sex-specific 5-year age-at-risk incidence rates of CVD in CPRD, among individuals without prior history of CVD, and not on statin treatment at baseline; 3) estimates for public health impact were shown, respectively, before and after recalibration.

**eTable 8: Numerical results for estimates of public health impact by additional assessment of polygenic risk score and clinical biochemistry markers, above conventional risk predictors, among 100,000 individuals**

| Conventional risk predictors | + PRS only | | | + Clinical biochemistry markers only | | | + Both | | |
|---|---|---|---|---|---|---|---|---|---|
| | 0-<5% | 5-<10% | ≥10% | 0-<5% | 5-<10% | ≥10% | 0-<5% | 5-<10% | ≥10% |
| **Cases (n=8219)*** | | | | | | | | | |
| 0-<5% | 899 | 194 | 2 | 856 | 229 | 9 | 872 | 210 | 13 |
| 5-<10% | 184 | 1372 | 446 | 168 | 1416 | 419 | 228 | 1226 | 548 |
| ≥10% | 0 | 211 | 3679 | 0 | 299 | 3591 | 1 | 442 | 3447 |
| **Non-cases (n=91,780)*** | | | | | | | | | |
| 0-<5% | 39,361 | 2809 | 6 | 39,299 | 2701 | 174 | 38,498 | 3454 | 224 |
| 5-<10% | 3956 | 16,944 | 2654 | 4146 | 16,864 | 2530 | 5955 | 14,134 | 3465 |
| ≥10% | 10 | 3037 | 15,666 | 6 | 3227 | 15,480 | 83 | 4647 | 13,982 |

* Among cases and non-cases, respectively, 1223 and 8570 participants had diabetes or LDL cholesterol measurement of 190 mg/dL or greater. Numbers in red are shown for individuals who were reclassified downwards with additional assessment, and numbers in blue are shown for individuals who were reclassified upwards with additional assessment.

**eFigure 1: Exclusion criteria applied in derivation of the primary analytic dataset**

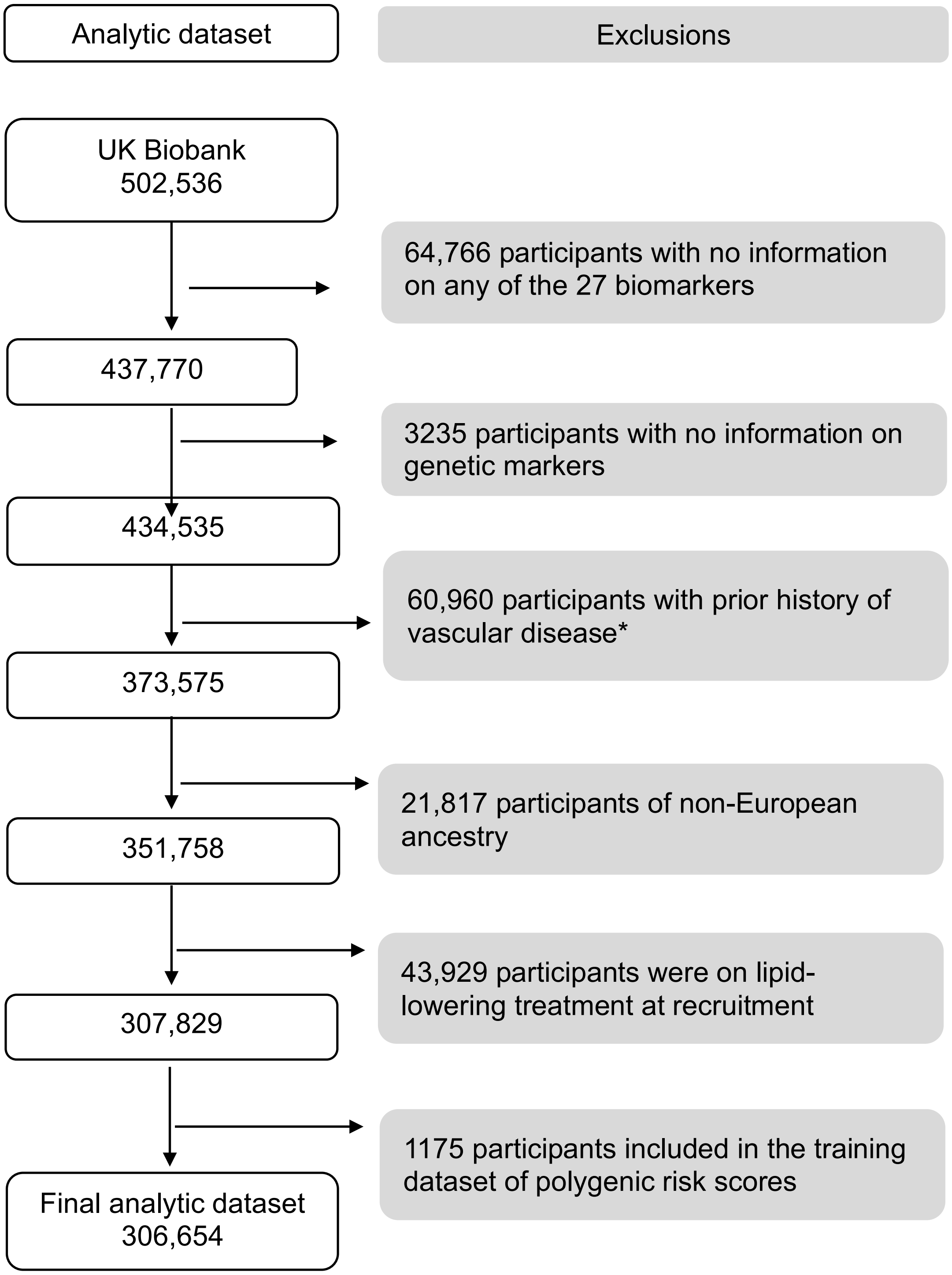


* Prior history of vascular disease at baseline included coronary heart disease, angina, other heart disease, stroke, transient ischaemic attack, peripheral arterial disease, and cardiac revascularisations.

**eFigure 2: Construction of polygenic risk score and selection of clinical biochemistry markers**

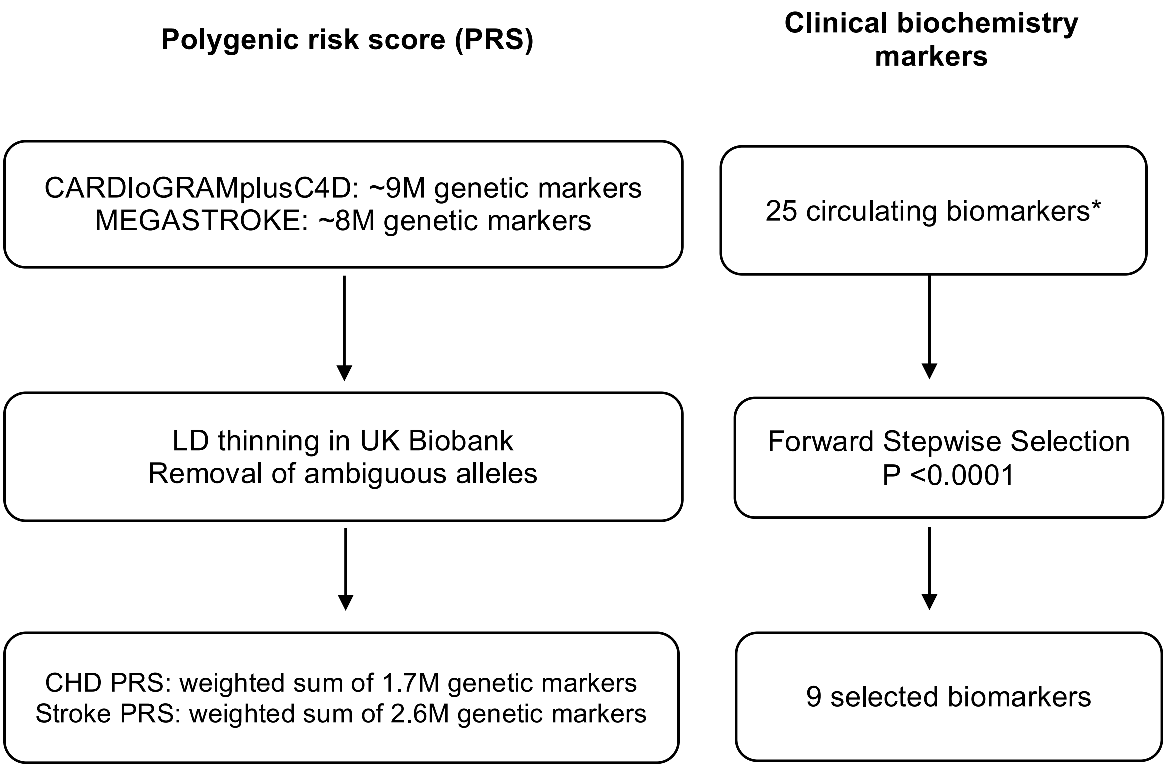


* Total cholesterol and HDL cholesterol were not included in the selection process because these two biomarkers were included in the reference model containing information on conventional risk predictors. LD, linkage disequilibrium; CHD, coronary heart disease; The numbers of single-nucleotide polymorphisms included were 1,743,179 in the PRS for CHD, and 2,595,401 in the PRS for stroke, respectively. The $r^2$ thinning threshold used was 0.9. Clinical biochemistry markers were selected using a forward stepwise procedure with Cox regression, stratified by centre and sex, adjusted for age at baseline, smoking status, history of diabetes, systolic blood pressure, total cholesterol and HDL-cholesterol levels. Selected biomarkers included Cystatin C, C-reactive protein, Lipoprotein (a), Sex hormone-binding globulin, Haemoglobin A1c, Creatinine, Albumin, Gamma-glutamyltransferase, Alanine transaminase, Alkaline phosphatase.

**eFigure 3: Process of estimating public health impact**

**Calculate 10-year CVD risk in UK Biobank using models including:**

- Conventional risk predictors*
- Conventional risk predictors + PRS only
- Conventional risk predictors + nine selected clinical biochemistry markers only
- Conventional risk predictors + both of the above

↓

**Recalibrate 10-year CVD risk for the above models, with age- and sex-specific CVD incidence rates from Clinical Practice Research Datalink**

↓

**Estimate reclassification statistics using recalibrated 10-year CVD risk, and risk categories defined by 2014 NICE guideline**

↓

**Apply reclassification estimates to 100,000 UK individuals with:**

- Age range: 40-75 years
- Age and sex profile as in standard UK population (2017)
- Age- and sex-specific CVD incidence rates as in Clinical Practice Research Datalink

* Conventional risk predictors included information on age, sex, smoking status, history of diabetes, total cholesterol and HDL cholesterol. Age- and sex-specific CVD incidence rates from the Clinical Practice Research Datalink were estimated among individuals without history of vascular diseases, and not on lipid-lowering treatment, at registration Records (1st Apr, 2004 – 21 Oct, 2017) on about 3.1 million patients were used in the present analysis

**eFigure 4: Shape and strength of associations of polygenic risk score with risk of CHD and stroke**

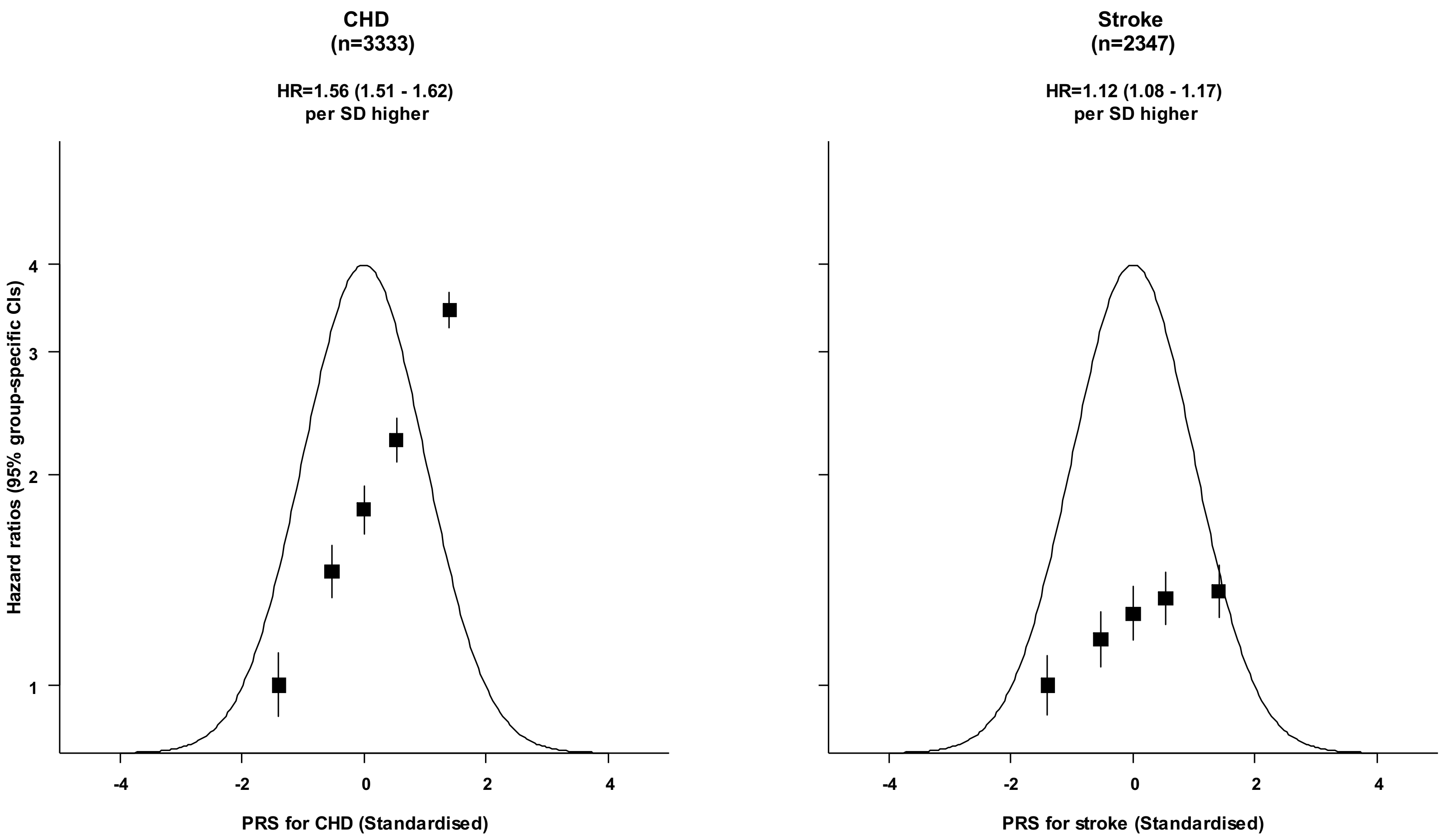


The shape of association was estimated by dividing all participants into fifths. Hazard ratios were estimated using Cox regression, stratified by study centre and sex, and adjusted for age at baseline. Each square has an area inversely proportional to the effective variance of the log risk in that specific group, with vertical lines representing the 95% confidence intervals.

**eFigure 5a: Shape of associations of polygenic risk score and clinical biochemistry markers with risk of CVD**

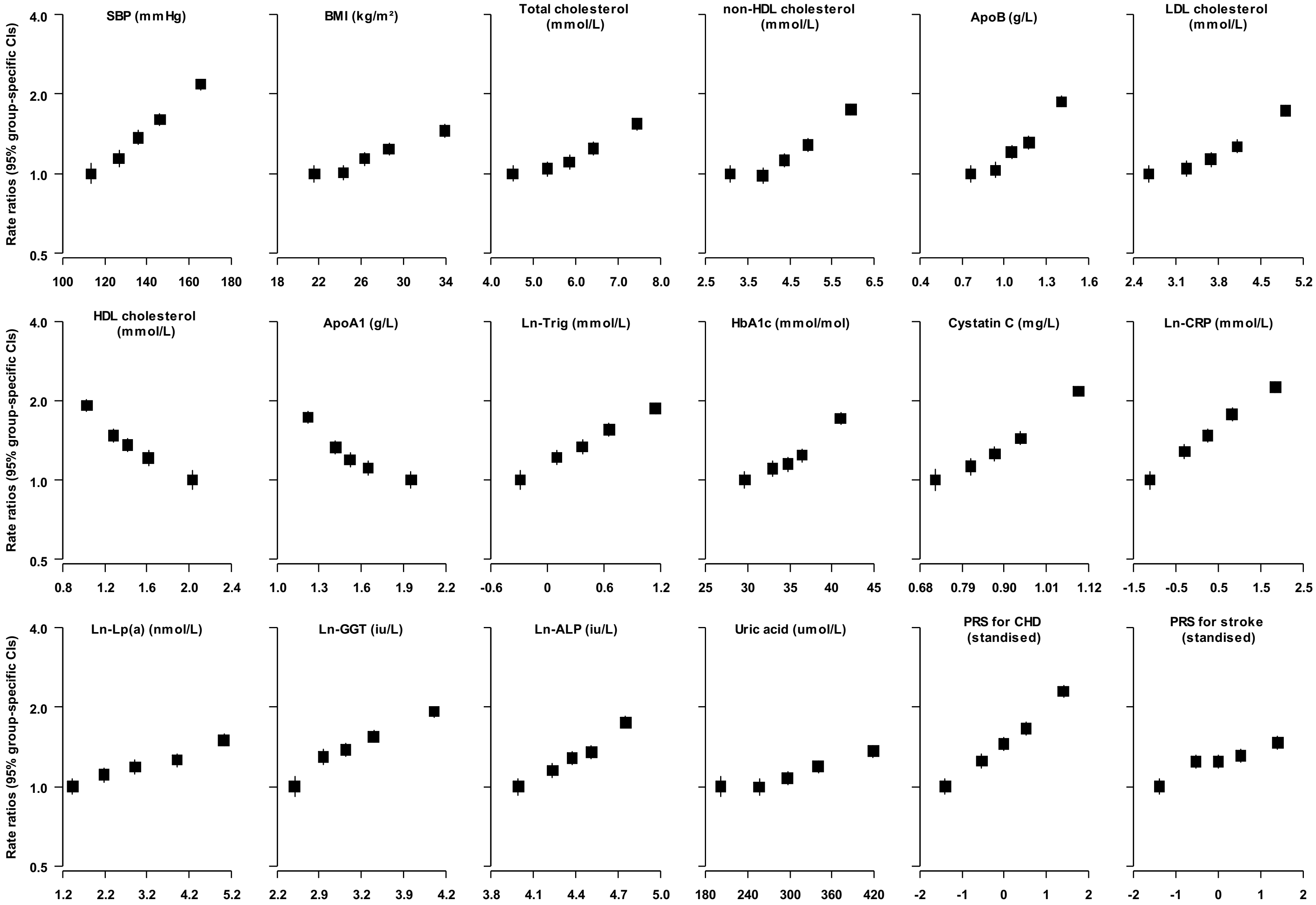


The shape of association was estimated by dividing all participants into fifths. Hazard ratios were estimated using Cox regression, stratified by study centre and sex, and adjusted for age at baseline. Each square has an area inversely proportional to the effective variance of the log risk in that specific group, with vertical lines representing the 95% confidence intervals.

**eFigure 5b: Shape of associations of polygenic risk score and clinical biochemistry markers with risk of CVD**

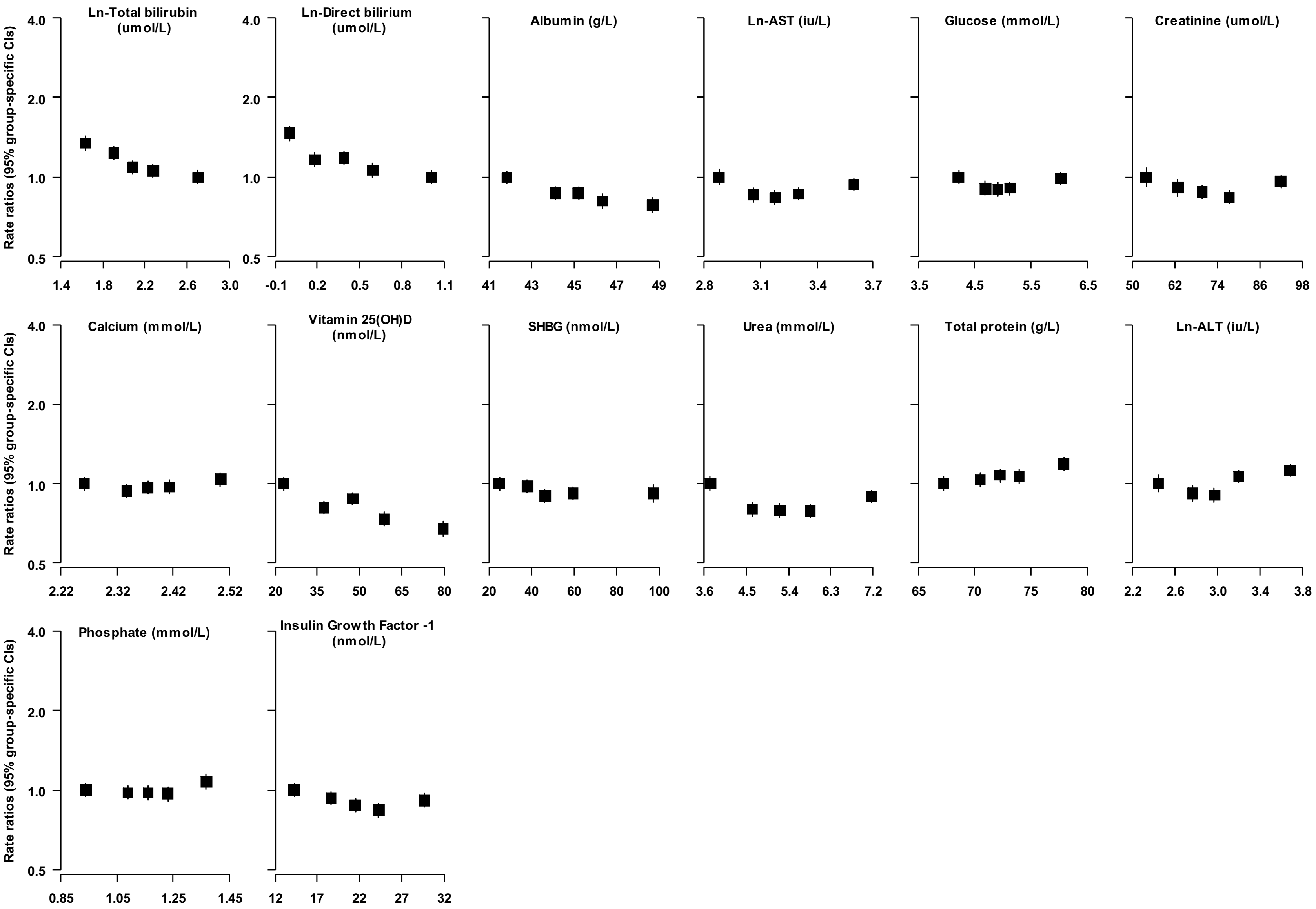


The shape of association was estimated by dividing all participants into fifths. Hazard ratios were estimated using Cox regression, stratified by study centre and sex, and adjusted for age at baseline. Each square has an area inversely proportional to the effective variance of the log risk in that specific group, with vertical lines representing the 95% confidence intervals.

**eFigure 6: Hazard ratios for CHD, stroke, and the composite CVD outcome, adjusted for conventional risk predictors**

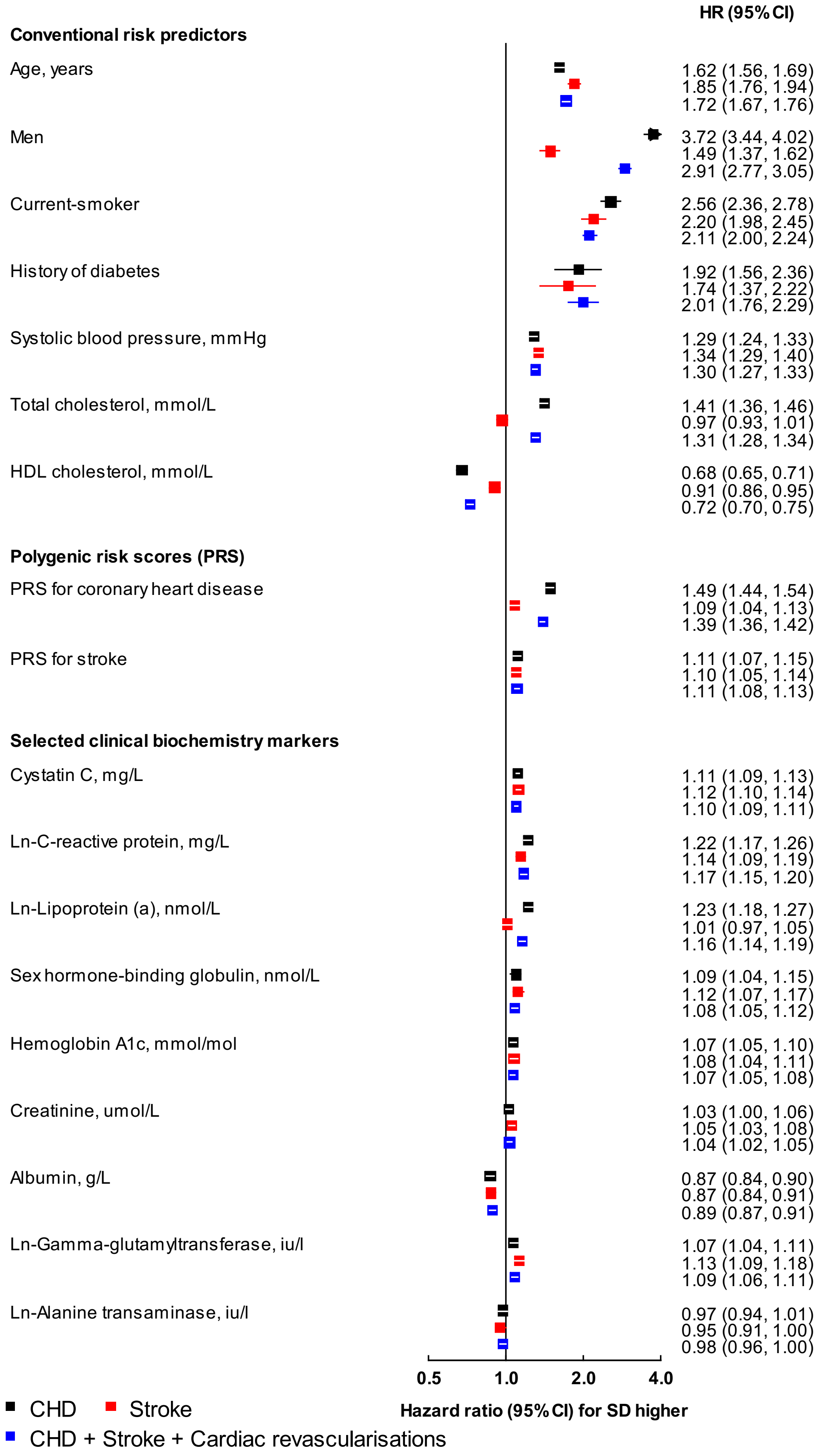


Hazard ratios (HRs) were estimated using Cox regression, stratified by study centre and sex, and adjusted for age at baseline, smoking status, history of diabetes, systolic blood pressure, total cholesterol and HDL-cholesterol levels, where appropriate. For continuous variables, HRs are shown for each SD higher of each predictor to facilitate comparison. For categorical variables, HRs are shown men vs. women, for patients with diabetes vs. without, for current smokers vs. others.

**eFigure 7: Incremental predictive ability of polygenic risk score and the nine selected clinical biochemistry markers for CVD outcomes, in isolation and in combination, above conventional risk predictors**

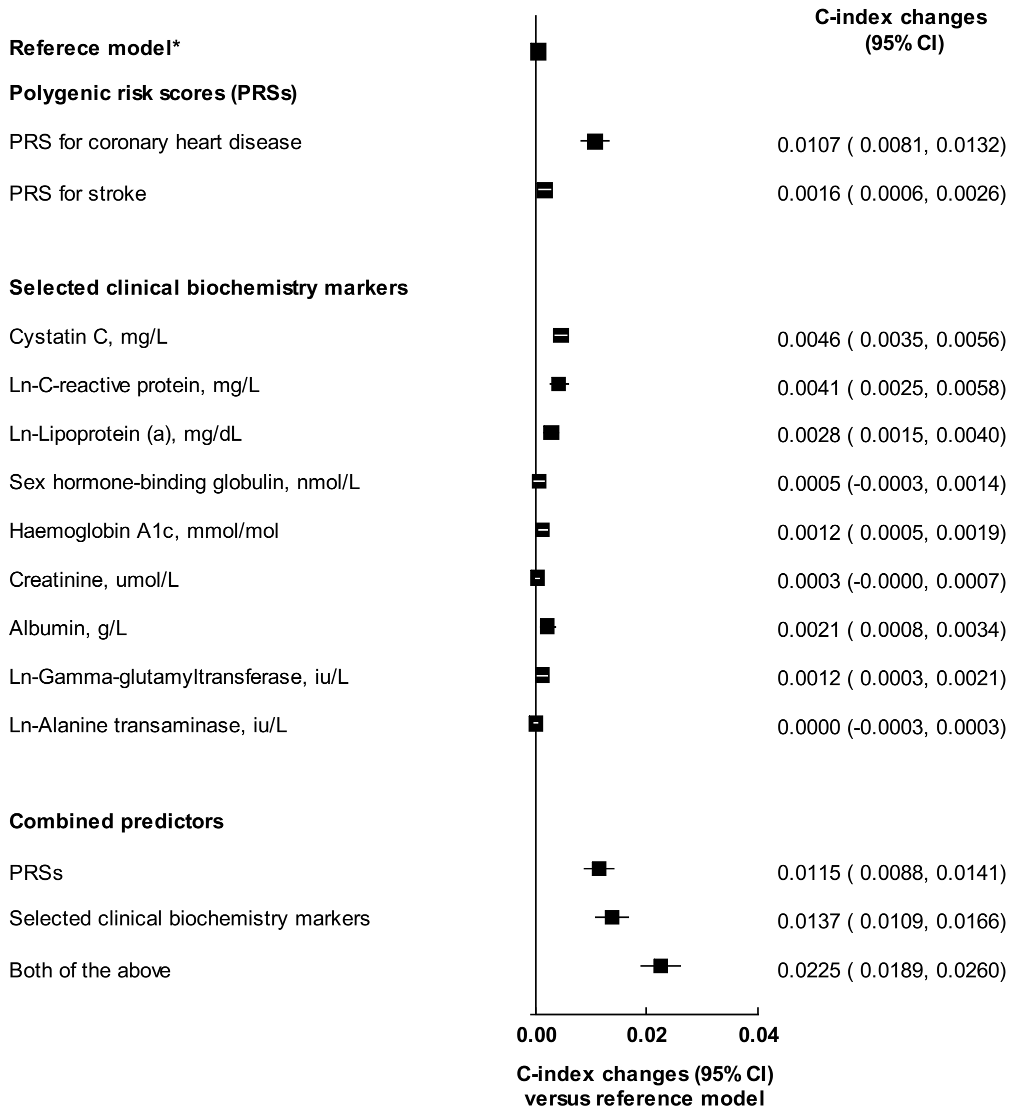


CVD, cardiovascular disease; *Reference model included information on age at baseline, sex, smoking status, history of diabetes, systolic blood pressure, total cholesterol and HDL-cholesterol levels.

**eFigure 8: Reclassification measures for CVD in UK Biobank by addition of polygenic risk score and clinical biochemistry markers, above conventional risk predictors**

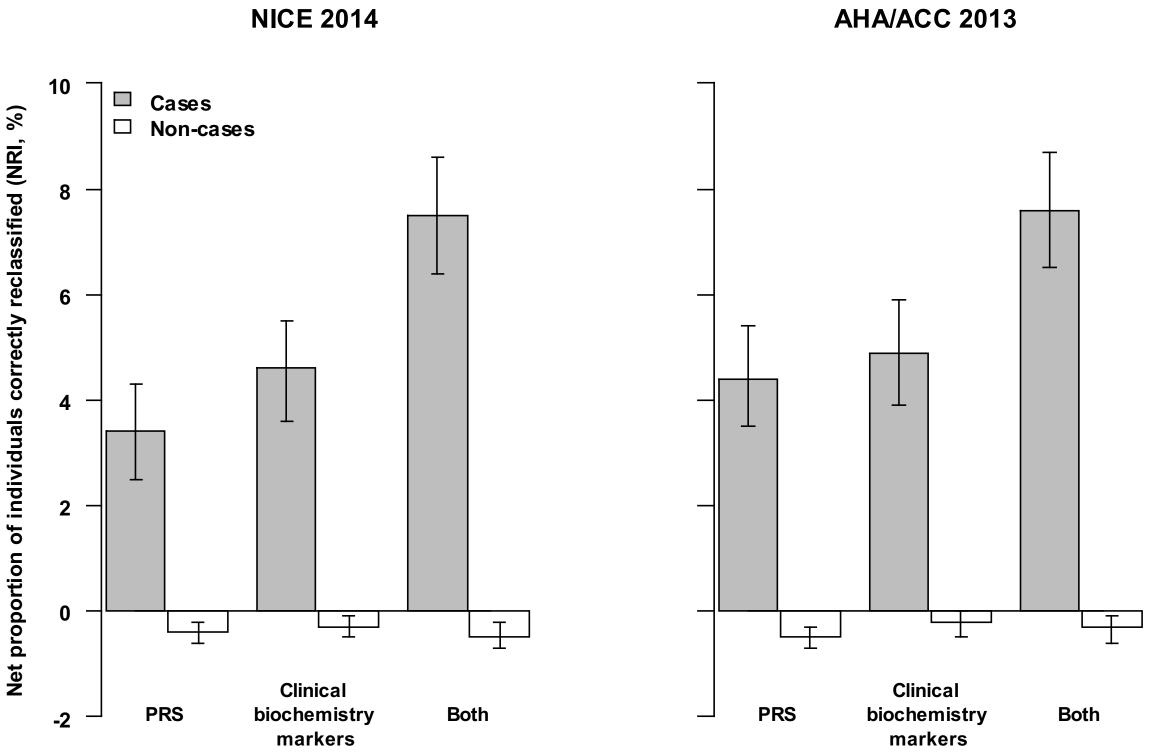


| | NICE 2014: PRS | NICE 2014: Clinical biochemistry markers | NICE 2014: Both | AHA/ACC 2013: PRS | AHA/ACC 2013: Clinical biochemistry markers | AHA/ACC 2013: Both |
|---|---|---|---|---|---|---|
| Cases | 3.4 (0.5) | 4.6 (0.5) | 7.5 (0.6) | 4.4 (0.5) | 4.9 (0.5) | 7.6 (0.6) |
| Non-cases | -0.4 (0.1) | -0.3 (0.1) | -0.5 (0.1) | -0.5 (0.1) | -0.2 (0.1) | -0.3 (0.1) |

ACC, American College of Cardiology; AHA, American Heart Association; NICE, National Institute for Health and Care Excellence; PRS, polygenic risk score; Conventional risk predictors included information on age at baseline, sex, smoking, systolic blood pressure, history of diabetes, total cholesterol and HDL-cholesterol levels. The predicted 10-year cardiovascular risk categories used to calculate the categorical NRIs were: <5%, 5-10%, ≥10%, according to the 2014 NICE guideline, and <5%, 5-7.5%, and ≥7.5%, according to the 2013 ACC/AHA guideline. Values of NRIs (s.e) for cases and non-cases in UKB are shown under each figure.

**eFigure 9: Incremental predictive values of polygenic risk score and clinical biochemistry markers, above conventional risk predictors, including body-mass index (BMI) or family history of CVD in the reference model**

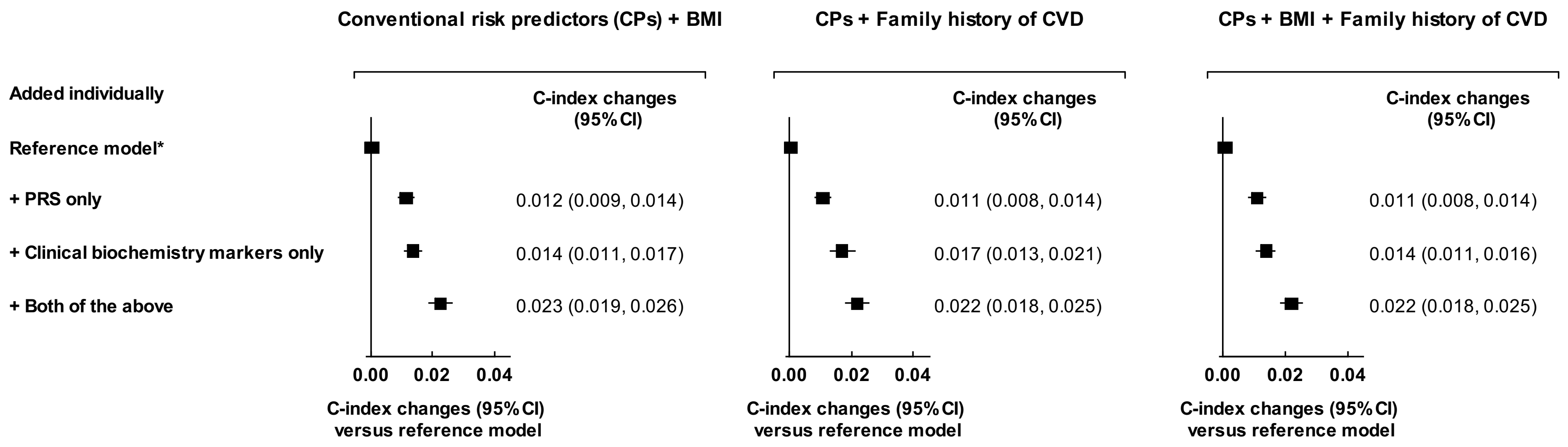


CVD, cardiovascular disease; CHD, coronary heart disease; PRS, polygenic risk score; *Reference model included information on age at baseline, sex, smoking status, history of diabetes, systolic blood pressure, total cholesterol and HDL-cholesterol levels. PRSs for CVD included PRS for CHD and PRS for stroke as two variables. Clinical biochemistry markers included Cystatin C, C-reactive protein, Lipoprotein (a), Sex hormone-binding globulin, Hemoglobin A1c, Creatinine, Albumin, Gamma-glutamyltransferase, Alanine transaminase.

**eFigure 10: Incremental predictive values of polygenic risk score and clinical biochemistry markers, above conventional risk predictors, with and without excluding participants on lipid-lowering treatment**

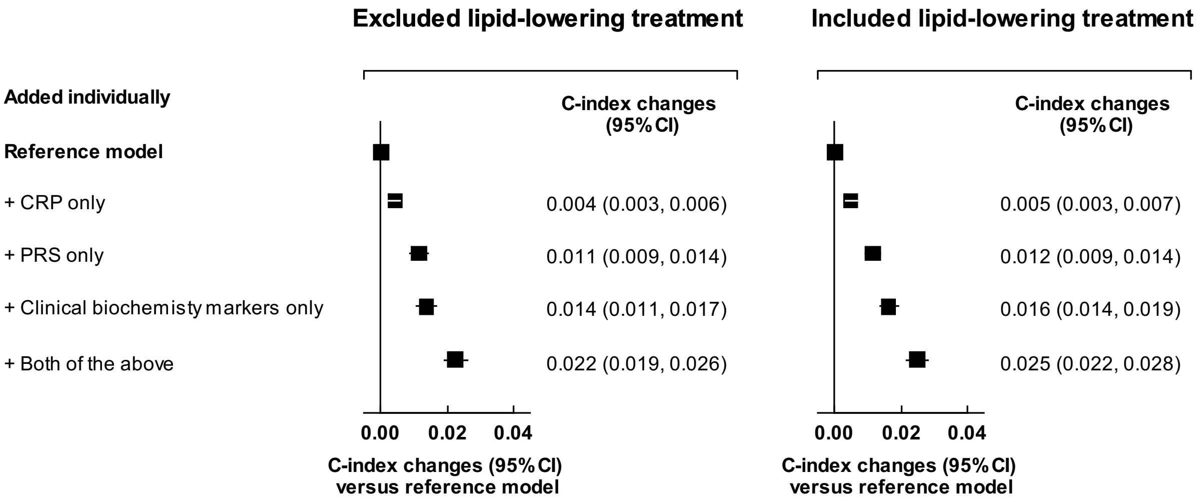


CVD, cardiovascular disease; CHD, coronary heart disease; PRS, polygenic risk score; *Reference model included information on age at baseline, sex, smoking status, history of diabetes, systolic blood pressure, total cholesterol and HDL-cholesterol levels. PRSs for CVD included PRS for CHD and PRS for stroke as two variables. Clinical biochemistry markers included Cystatin C, C-reactive protein, Lipoprotein (a), Sex hormone-binding globulin, Hemoglobin A1c, Creatinine, Albumin, Gamma-glutamyltransferase, Alanine transaminase.

**eFigure 11: Incremental predictive values of polygenic risk score and clinical biochemistry markers, above conventional risk predictors, for CVD outcomes, with and without including revascularisation procedures**

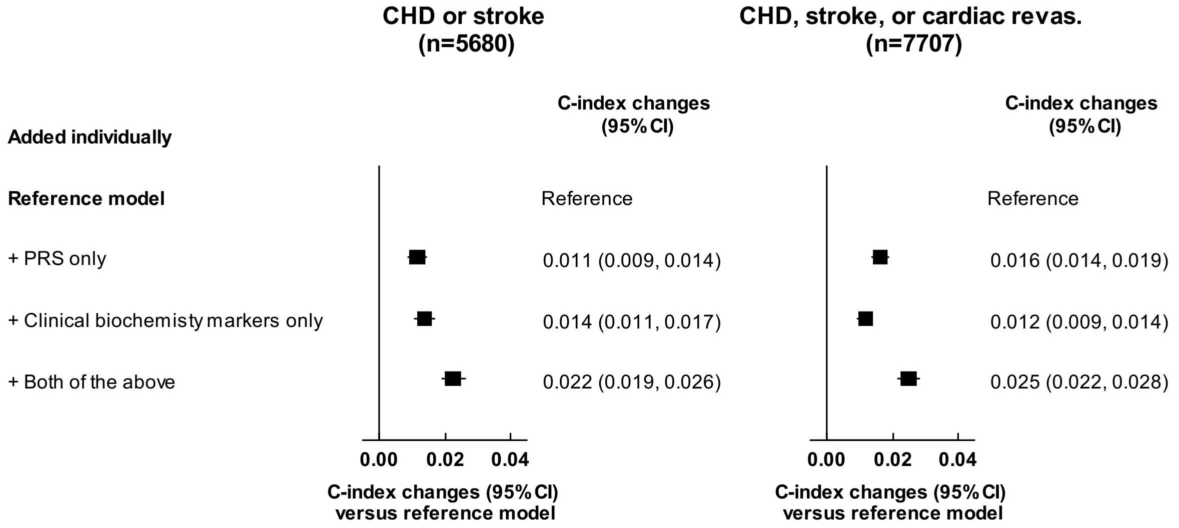


CVD, cardiovascular disease; CHD, coronary heart disease; PRS, polygenic risk score; Reference model included information on age at baseline, sex, smoking status, history of diabetes, systolic blood pressure, total cholesterol and HDL-cholesterol levels. PRSs for CVD included PRS for CHD and PRS for stroke as two variables. Clinical biochemistry markers included Cystatin C, C-reactive protein, Lipoprotein (a), Sex hormone-binding globulin, Hemoglobin A1c, Creatinine, Albumin, Gamma-glutamyltransferase, Alanine transaminase. Left panel includes myocardial infarction (I61-63), and fatal coronary heart disease (I64-65). Right panel also includes revascularisation procedures (OPCS-4: K40-K46, K49, K50.1, K50.2, K50.4, or K75).

**eFigure 12: Estimates of public health impact with targeted assessment of polygenic risk score, and clinical biochemistry markers among 100,000 UK adults using AHA/ACC guideline**

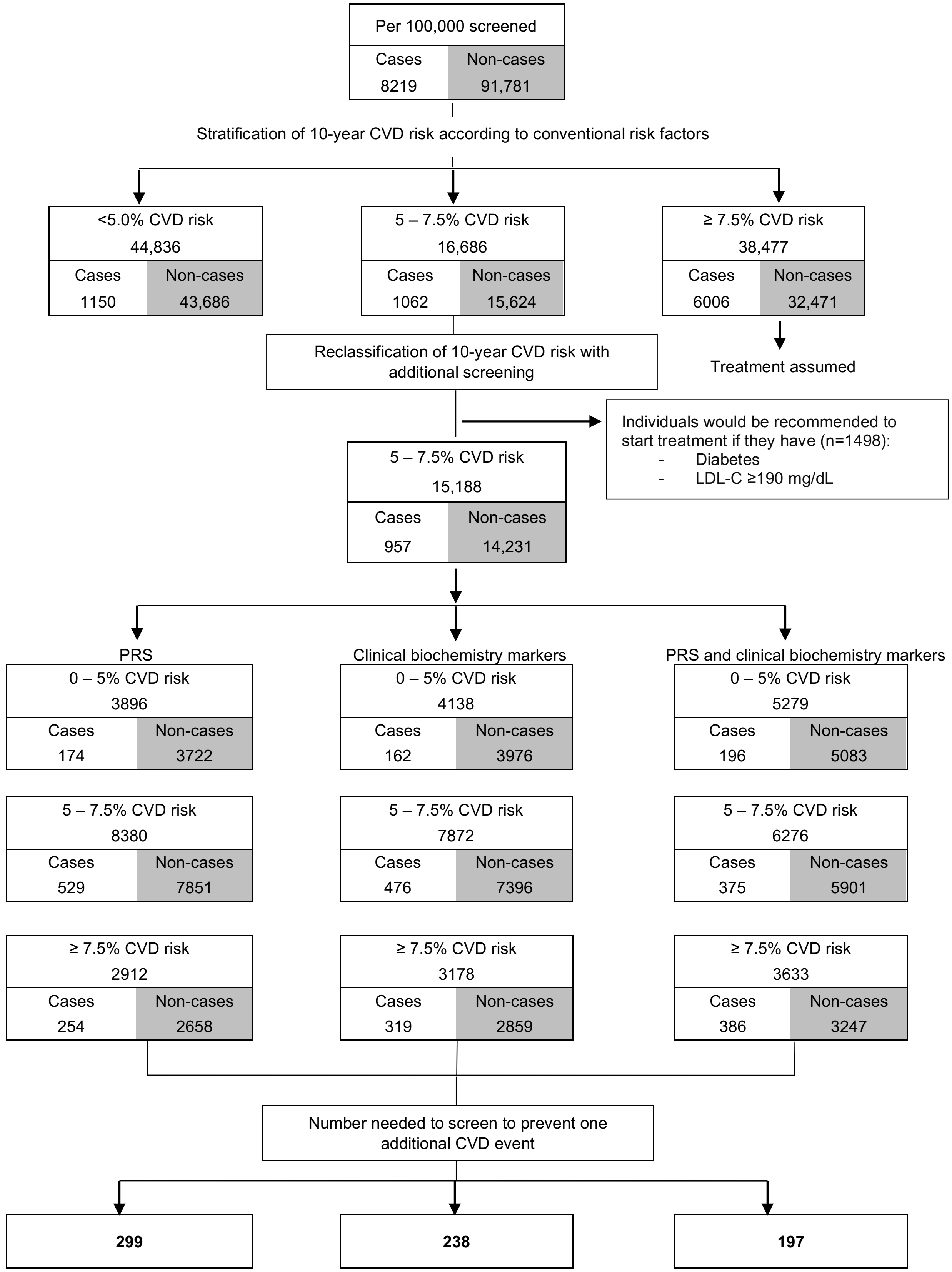

**eFigure 13: Estimates of public health impact by additional assessment of polygenic risk score and clinical biochemistry markers, above conventional risk predictors, among 100,000 individuals**

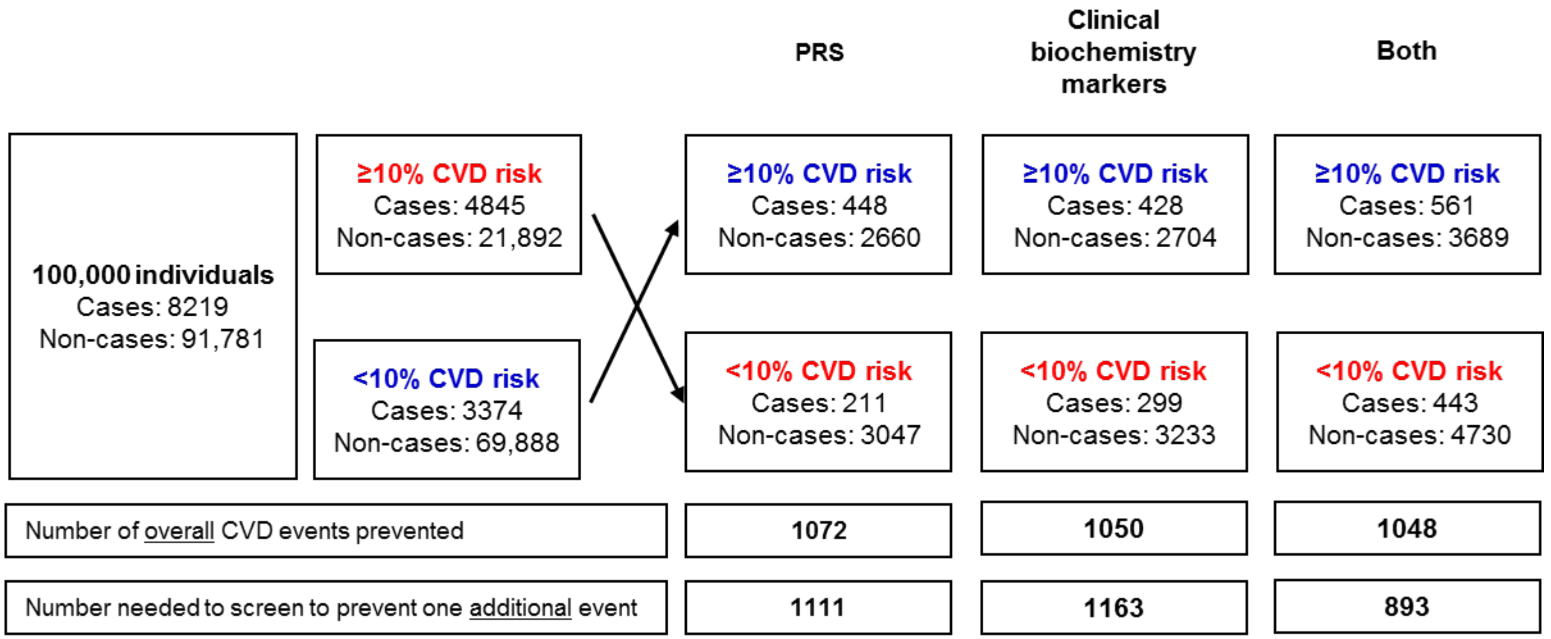


Numbers in red are shown for individuals who were initially at high-risk and were reclassified down to intermediate-risk; numbers in blue are shown for individuals moving from intermediate-risk to high-risk. Among 100,000 individuals, 1233 cases and 7338 non-cases were treated, irrespective of their 10-year CVD risk, since they had history of diabetes, or LDL cholesterol ≥190 mg/dL. Number of cases screened as high-risk or due to diabetes, LDL cholesterol levels, using conventional risk predictors alone were 5122, and thus, events prevented were 1024 (5122 × 0.2).

13. National Institute for Health and Care Excelllence. Cardiovascular disease: risk assessment and reduction, including lipid modification. London: National Institute for Health and Care Excellence 2014.
